# FLAG-X: Hybrid machine learning workflows for automated gating of clinical flow cytometry data

**DOI:** 10.64898/2026.01.10.698765

**Authors:** Paul Martini, Marziyeh Mohammadi, Michael C. Thrun, David B. Blumenthal, Stefan W. Krause

## Abstract

Flow cytometry analysis is widespread practice in cell biology, immunology and hematology. Cell populations of interest are typically identified by consecutively examining the expression levels of antigen marker pairs. Since this manual gating process lacks standardization and is time-consuming, several machine learning (ML) methods for automated gating of flow cytometry data have been proposed in recent years. However, their translation into routine workflows has been limited. To address this, we developed the Python package FLAG-X (“**fl**ow cytometry **a**utomated **g**ating toolbo**x**”), which supports two novel workflows that integrate manual with ML-based gating, using labeled and unlabeled training data. We selected state-of-the-art ML methods developed for automated gating for inclusion in FLAG-X, based on their gating performance in comparison to manual expert annotations. FLAG-X provides a unified interface for top-performing methods and enables seamless integration with standard software for manual gating by exporting results as FCS files. To demonstrate its practical utility, we applied FLAG-X to representative cases from clinical practice. FLAG-X is available at https://anaconda.org/channels/bioconda/packages/flagx/overview.

**Author summary:** In our research, we work with flow cytometry data, a common laboratory technique to measure the expression of specific antigens in individual cells of a patient sample. In everyday clinical diagnostics or research, experts use dedicated software tools to “manually” inspect these data and often select a subset of cells for further analysis, e.g. B cells for Lymphoma subtype classification. However, this manual procedure, known as gating, can be time-consuming and is not standardized such that outcomes may vary based on the clinician carrying out the cell selection. Automated, machine learning-based methods developed to mitigate those problems are rarely used in routine clinical work. In this study, we set out to close this gap by developing novel workflows that allow clinicians to more easily integrate (semi-)automated approaches into their existing workflows. To do this, we we developed a software package that brings together top-performing methods, functionality to handle clinical flow cytometry data and is designed to be compatible with with existing tools for manual analysis. We evaluated our workflows and toolbox using realistic clinical scenarios and considered practical requirements for their use. By integrating automation into familiar workflows, we strive to make flow cytometry gating both faster and more consistent to support wider use of computational methods in clinical diagnostics.

## Introduction

Flow cytometry (FCM) is a laboratory technique that is widely used for analyzing the physical and chemical characteristics of cell suspensions at single-cell resolution in a high-throughput fashion. Besides its broad application in research, FCM has become standard clinical practice for analyses and diagnosis in hematology and immunology. FCM data is typically represented as a matrix *X* = (*x*_*i,j*_) ∈ ℝ^*n×m*^, where each of the *n* rows corresponds to a single event (i.e., a cell) and each of the *m* columns corresponds to a parameter measured by the quantity of emitted light after excitation with laser beams. These parameters consist of light reflected in the forward and orthogonal direction (commonly referred to as forward and side scatter), as well as fluorescent light emitted by cells binding fluorochrome-labeled antibodies or other fluorescent dyes.

Manual FCM analysis is a multistep process [1]. First steps are check of data integrity, elimination of artifacts and correction of fluorochome spillover. The next crucial step is the definition of cell populations of interest, typically performed by consecutively drawing boundaries (gates) around clusters of cells in two-dimensional (2D) scatter plots of suitable marker pairs. Populations are then quantified either as an end result (e. g., percentage of CD3+ CD4+ T lymphocytes) or used as starting point for higher level analysis (e.g. leukemia diagnosis). However, this procedure is inherently subjective, time-consuming, and difficult to standardize [2]. Notably, [3] reported that inter-rater reliability among experts performing manual gating can be as low as *κ* = 0.42 (Cohen’s kappa, *κ* = 0 denotes no agreement and *κ* = 1 denotes complete agreement) depending on the cell type to be analyzed. While dedicated protocols for standardization of manual gating were proposed for selected scenarios [4], they are not available for the majority of practical tasks. Also, analyzing only two markers at a time may overlook cell populations that are only distinguishable when considering a higher-dimensional space. These limitations have motivated the development of automated, data-driven approaches that aim to provide objective, scalable, and standardized means of analyses.

According to recent citation trends, the most popular methods for automated FCM data analyses are unsupervised approaches [5]. Indeed, numerous clustering algorithms have been developed for identifying known cell populations as well as delineating novel subsets during exploratory analyses [6–11]. These methods have been comprehensively evaluated and some of them shown to align quite well with expert manual gating [3, 12]. Among these unsupervised methods, FlowSOM [10], a method based on self-organizing maps (SOM) [13], stands out as it consistently ranks as one of the top-performing tools [3, 12] and has been applied across a wide range of published studies [14]. However, unsupervised methods alone are of interest for hypotheses generation in research, but are insufficient for many other scenarios for several reasons. First, the resulting cell populations still require manual annotation. Second, the number of distinct populations within a sample must either be predefined or inferred automatically, which can be unreliable. Third, different unsupervised methods utilize a wide range of alternative mathematical frameworks, which typically produce equally valid, but divergent partitions of events. This leads to inconsistencies in population delineation limiting comparability and reproducibility across tools and studies. Finally, in many cases, only a minority of the formally distinguishable populations within a sample are of biological or medical interest.

To achieve end-to-end automation, supervised methods that directly assign cell-level annotations are essential. Supervised gating methods can broadly be categorized into classical machine learning (ML) approaches such as GateMeClass [15], linear discriminant analysis [16], bayesian trees [17], and ACDC [18] and on the other hand deep learning-based techniques such as DGCyTOF [19] and DeepCyTOF [20]. However, a key limitation of existing ML-based automated gating methods is that they do not address the requirements for deployment in conventional FCM workflows. In essence, existing approaches are supervised classifiers that require FCM data with ground-truth cell type annotations for training. In practice, such annotated training data are rarely available in clinical settings. For clinical translation, automated methods must therefore be embedded into workflows that integrate with standard manual gating procedure, support the generation of high-quality annotated training data, and provide mechanisms to identify and correct annotation inconsistencies. Furthermore, such an automated gating tool should operate on all collected events without prior exclusion of low-quality events and be trainable with as few annotated datasets and events as possible to run on a conventional computers commonly available in clinical environments.

This motivated us to develop the Python package FLAG-X (“**fl**ow cytometry **a**utomated **g**ating toolbo**x**”). FLAG-X addresses the limitations of existing ML-based gating methods through two complementary workflows for unlabeled and labeled input data, respectively. The first workflow leverages learned representations of the training data as an aid for manual gating and supports semi-automated gating of novel samples through mapping of cells to SOM nodes and visual inspection of 2D representation spaces with standard gating software. The second workflow can be used to annotate samples via fully automated ML-based gating methods trained on annotated FCM data (cell type annotations can be obtained using the the first workflow), while incorporating an expert-in-the-loop mechanism to identify and correct unreliable “ground truth” training annotations. FLAG-X provides a unified interface for top-performing ML-based gating methods, implements comprehensive functionality for FCM data processing in Python, and supports import and export of Flow Cytometry Standard (FCS) files for seamless integration with standard manual gating software to facilitate reproducible, interpretable, and scalable FCM data analysis. FLAG-X is available as an open-source Python package at https://anaconda.org/channels/bioconda/packages/flagx/overview.

## Materials and methods

### Machine learning models for automated gating

To select ML-based gating methods for inclusion in FLAG-X, we evaluated the ability of state-of-the-art approaches to reproduce expert-provided gating, which was treated as ground-truth. We included (1) GateMeClass [15], a recent classical ML approach based on Gaussian mixture models (GMMs), (2) DGCyTOF [19], a state-of-the-art deep learning model that combines a multilayer perceptron (MLP) classifier with correlation-based calibration, (3) DGCyTOF’s core MLP classifier without correlation-based calibration, and (4) SOM classifier, a supervised variant of a SOM implemented by ourselves, which we included in the comparison due to the popularity of FlowSOM.

#### GateMeClass

GateMeClass relies on GMMs to discretize per-cell marker expression into categorical levels (low, medium, high). To account for skewed marker distributions, it employs ranked set sampling before GMM fitting. During the training phase, the method constructs a marker table that encodes decision rules based on the expression level of the most discriminative marker for each pair of cell types. These discriminative markers are identified by their feature importance for classification in a tree-based model. For annotation, each cell’s marker expression profile is mapped to this marker table to assign an initial label. Cells that remain unassigned after this mapping are further refined using a *k*-nearest neighbors classifier, which assigns final labels based on local similarity in marker space.

#### DGCyTOF

DGCyTOF uses MLP classifier to predict cell type probabilities. During prediction, cells with low-confidence predictions are reclassified in a correlation-based calibration step. A sensible confidence threshold is computed on a validation set which is held out from model training.

#### MLP classifier

The MLP classifier consists of a fully connected neural network with ReLU [21] activation in the hidden layers and softmax activation in the output layer. Three layers with 128, 64, and 32 hidden units, respectively, were used.

#### SOM classifier

The SOM classifier builds on a SOM which is trained in an unsupervised manner to project high-dimensional marker expression data onto a two-dimensional grid of units, preserving the topological structure of the input space. Following this first part of the training phase, each unit on the grid is assigned a cell type label through majority voting based on the labeled training data. During inference, new cells are mapped to their best matching unit in the SOM grid and annotated according to that unit’s label. The cell type fractions at the best matching unit can be interpreted as a probabilistic prediction. Technical details are provided in Section S1 in S1 Appendix.

### Datasets and preprocessing

The candidate ML-based gating methods described above were tested on four different expert-annotated datasets: Flowcyt [22], Imstat, LT1, and LT2. Moreover, we used binary classification variants of the LT1 and LT2 datasets, denoted as LT1b and LT2b, in which subpopulations were aggregated into broader classes. No manual preselection of events was performed, that is, we used the datasets as-is without removing debris. Compensated data (max. = 2^20^) was used directly. For the private datasets (Imstat, LT1, LT2), we applied a log_10_-transformation with biologically informed lower cut-offs defined per channel. For the Flowcyt dataset, we applied the same transformation but with a uniform cut-off of 100 across all channels. Input data for GateMeClass was arcsinh transformed with a co-factor of 150 since lower-bound transformations resulted in stability issues described in Section S2.1 in S1 Appendix. Further details on the individual datasets are provided in the following paragraphs. Dataset sizes, class distributions, ground-truth gating information, and mapping of letter labels to biological cell types are provided in Figures S1–S2 in S1 Appendix and in Tables S2–S5 in S1 Appendix.

#### Flowcyt

Flowcyt is a dataset of healthy bone marrow samples and was curated specifically for multi-class classification benchmarking in FCM. We used the cell type annotations provided by the authors for model training and evaluation.

#### Imstat

Imstat is a dataset of immune status analyses performed with different lysis reagents, which leads to additional differences between the samples on top of the expected biological variability [23]. Six cell populations plus one class of remaining cells and one class of irrelevant events were defined using forward scatter, side scatter, and 8 antibody marker channels. Several cell populations are clear-cut while others are more difficult to distinguish from the rest of the cells [24]. Manual cell type annotation was performed by one expert operator at the University Hospital of Erlangen.

#### LT1, LT2 and LT1b, LT2b

The lymphoma tubes datasets LT1 and LT2 contain measurements from patients with different B cell non-Hodgkin’s lymphoma, chronic lymphatic leukemia - CLL, mantle cell lymphoma - MCL, follicular lymphoma - FL, marginal zone lymphoma - MZL, lymphoplasmacytic lymphoma - LPL, hairy cell leukemia - HCL, diffuse large B cell lymphoma - DLBCL, Burkitt lymphoma - BL, monoclonal B cell lymphocytosis - MBL, unclassifiable B cell lymphoma - UC, and normal controls (NB), a total of 99 cases, preselected from a larger number of samples to evenly cover a broad range of diagnoses, collected at the University Hospital of Marburg, Germany [25]. Because the phenotypes of the lymphomas differ substantially between samples, gating on the LT1 and LT2 datasets is expected to be a more difficult task than gating on the Flowcyt and Imstat datasets. Each patient is represented by two complementary FCS files, each corresponding to a tube with a specific marker panel. A special feature of these measurements is that more than one antibody was used in some of the channels (see Table S6 in S1 Appendix), which makes the analysis more complicated. For our benchmark, the data were used in two ways: For a more detailed analysis, “healthy” B cells, dying B cells (defined by low forward scatter), CD45-negative events (“debris”, LT1 only), and the rest of the cells were distinguished. For a dichotomous evaluation (LT1b and LT2b), B cells including dying B cells were lumped together and separated from all other events (Tables S2 and S3 in S1 Appendix). Manual cell type annotation was performed by one expert operator at the University Hospital of Erlangen.

### Testing of machine learning models

In an initial round of tests, gating performance was evaluated for all tested models across all datasets to select top-performing models for inclusion in FLAG-X. In addition, we analyzed FLAG-X’s training data and computational resource requirements, as well as precision-recall trade-off of the included methods in the context of a cell selection task prior to downstream analyses.

#### Data splitting

Each dataset was randomly split into training and test sets using a 60:40 ratio at the sample level. For the LT1, LT2, LT1b, and LT2b datasets, the splits were stratified with respect to lymphoma subtypes. The data matrices of the training samples were concatenated and shuffled along the event dimension before training. Antibody marker channels on which models were trained are shown in Table S6 in S1 Appendix.

#### Hyperparameter selection

For training the GateMeClass, DGCyTOF, and MLP models, we used the default parameters provided by [15] and [19], respectively. For SOM classifier, we conducted a nested grid search on a reduced version of the Imstat training set, obtained by concatenating all training samples and downsampling to 10% of total events. While this gives SOM classifier an advantage on the Imstat dataset, the comparison remains equitable for all other datasets. In stage one, each hyperparameter was varied individually over a broad range while all others were fixed to Somoclu’s [26] defaults. Performance was evaluated by macro F1 scores using 3-fold cross validation. In stage two, we performed an extensive grid search over the most promising parameter values, splitting the reduced training set into 66% for training and 34% for validation to reduce runtime (performance was observed to be stable across folds). The number of epochs was initially set high to ensure stable validation performance and then reduced to a smaller value that still produced stable results for the optimal parameter combination. A description of the SOM’s hyperparameters and their influence on model training, based on the first tuning stage, is provided in Table S7 in S1 Appendix. The hyperparameter grid of the second stage and the selected optimal values are given in Table S8 in S1 Appendix. Note that for training with reduced training datasets on local machines, we used SOM classifier with a grid of size (20,20) rather than the (25,25) grid identified as optimal during hyperparameter tuning. This smaller grid was sufficient for the reduced training data and avoids oversparsification of the SOM.

#### Performance score computation

Performance scores were computed per test sample and then averaged. For classes *i ∈ {*1, …, *k}* and test samples *j ∈ {*1, …, *l}* the macro F1 scores are computed as

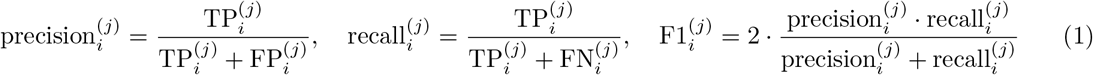

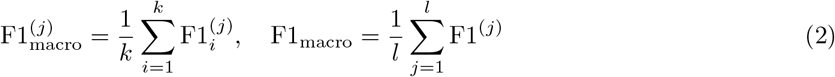

with 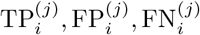 being the true positives, false positives, and false negatives for class *i* and sample *j*. Given the set of performance scores 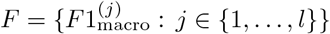, empirical confidence intervals were estimated using a nonparametric bootstrapping approach. Specifically, *B∈* ℕ bootstrap samples 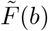, *b* = 1, …, *B*, were generated by resampling the original set *F* with replacement. For each bootstrap sample, the average macro F1 score was computed according to Eq (2), resulting in the bootstrap estimates 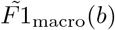 for *b* = 1, …, *B*. Finally, the 1 *− α, α ∈* (0, 1) confidence interval was determined by taking the *α/*2 and 1 *− α/*2 empirical quantiles of the distribution of the bootstrap estimates.

#### Model testing with limited training data

To evaluate the feasibility of training when only few labeled samples are available for training, we assessed the performance of the ML models selected for FLAG-X on subsets of the training datasets. For this, samples were incrementally added in randomized order to form progressively larger training sets. This way, each training set was a subset of the next larger one until all samples were included. For downsampling the number of events per sample, we applied a sample-wise stratified strategy, reducing each sample to a target number of events while preserving the within-sample proportions of cell types.

### Implementation

For GateMeClass and DGCyTOF, we used the R and Python implementations provided by their respective publications. The implementation of MLP was adapted from the DGCyTOF source code (https://github.com/lijcheng12/DGCyTOF/, accessed on August 12, 2025). Our Python implementation of SOM classifier uses Somoclu [26], a C++ based, parallelized package for training SOMs that includes a Python interface. This ensures scalability and fast training, making it feasible to train large SOMs (grid dimensions (*≥*10, 10)) on large datasets (number of events *≥*15,000,000).

The FLAG-X Python package consists of three core modules: io for loading FCM data in CSV format, FCS format (supported versions: 2.0, 3.0, and 3.1), or LMD format (containing an FCS2.0, FCS3.0, or FCS3.1 compliant part) and saving results back to FCS, `dimred` for computing dimensionality reductions, and gating containing implementations of MLP and SOM classifier. To enhance usability and enable seamless integration into existing ML workflows, the gating methods in FLAG-X conform to scikit-learn’s [27] estimator API, providing the familiar `.fit()` and `.predict()` methods for model training and inference. Additionally, FLAG-X offers higher-level access via an end-to-end pipeline that can be executed through a command-line interface.

### Computational environment

Benchmark runs on the full datasets were carried out on the Woody and TinyGPU high-performance computing (HPC) clusters hosted at the Erlangen National High Performance Computing Center (NHR@FAU, see Table S9 in S1 Appendix for details). Additional benchmark runs of MLP and SOM classifier on a local machine were performed on a Lenovo ThinkPad P14s Gen 5, equipped with an Intel Core Ultra 7 165H processor, an NVIDIA RTX 500 Ada Generation Laptop GPU with 4 GB of VRAM, and with 8 CPU cores allocated.

## Results

### Three out of four tested methods reach useful gating performance

Fig 1 presents macro F1 scores obtained in our gating benchmark, computed as the unweighted average of class-wise F1 scores, for each sample across datasets. GateMeClass shows substantially lower performance compared to the other methods and exhibits high variability across samples. We observed no clear performance gain in using DGCyTOF over its underlying MLP. Both DGCyTOF and MLP classifier achieve consistently high performance, with median macro F1 scores generally exceeding 0.9 and low variance across samples. SOM classifier performs comparably to DGCyTOF and the MLP on the Imstat, LT1b and LT2b datasets but underperforms on datasets with extreme class imbalance, where minority classes are heavily underrepresented. Particularly, the prediction of difficult-to-gate classes such as dying B cells in LT1 and LT2, and HSPCs and B cells in Flowcyt was found to be challenging. Interestingly, debris and other events outside the leukocyte region (as represented by population X in the Imstat dataset), that are usually sorted out in the first phase of manual gating, were detected with high reliability. Thus, prior “pre-cleaning” of raw data is not necessary using these classifiers. A detailed analysis of class-wise prediction performance is provided in Fig S3 in S1 Appendix. Additionally, the concordance between predicted and true cell type proportions for each test sample of each dataset is shown in Fig S4 in S1 Appendix, and a qualitative assessment of the methods is provided in Section S2 in S1 Appendix. Overall, three of the four evaluated methods (DGCyTOF, MLP classifier, and SOM classifier) achieved gating performances that meet clinically useful standards across multiple datasets, with limitations primarily in the presence of difficult-to-gate minority classes.

**Fig 1.**
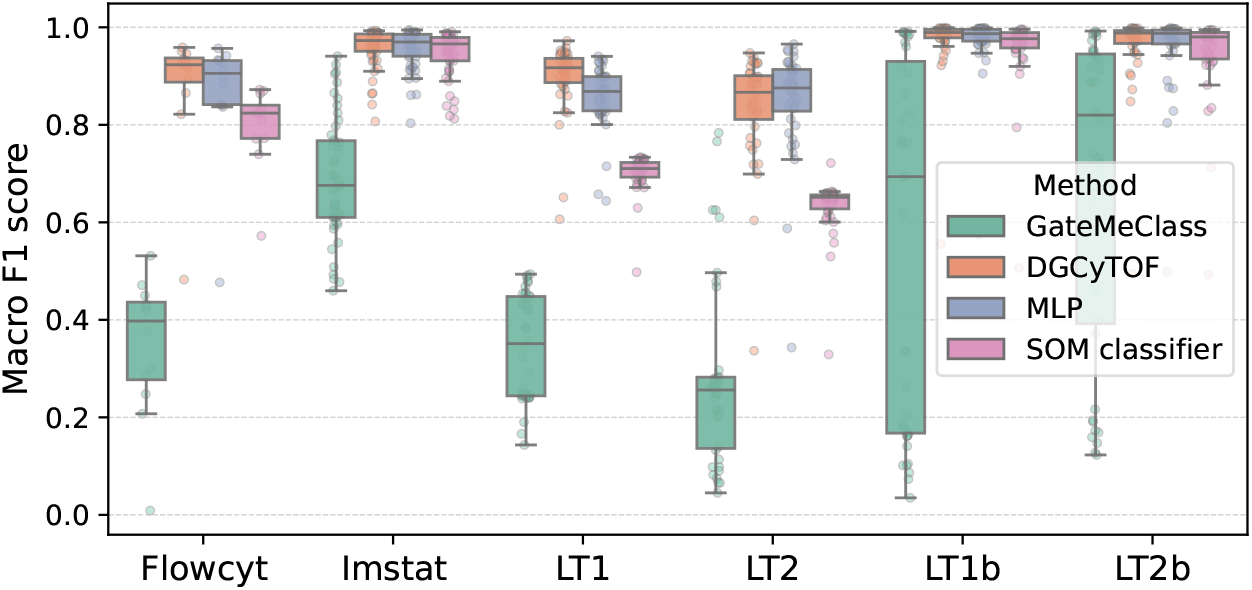
Gating performance of tested methods. Box plot of sample-wise macro F1 scores across datasets for each method.

### FLAG-X introduces novel workflows to integrate automated and manual gating

In our FLAG-X Python package, we implemented the best-performing methods from our benchmark: the MLP and the SOM classifier (DGyCyTOF was not selected because it does not improve upon its core MLP classifier). FLAG-X provides a unified interface for both approaches, ensures compatibility of input and output data with standard FCM analysis software, and supports two expert-in-the-loop workflows that interlock automated gating with manual gating, inspection, and refinement and are applicable in two different application scenarios (Fig 2, Jupyter notebooks that show how to run the two workflows are available in FLAG-X’s GitHub repository (https://github.com/bionetslab/FLAG-X):

- Application scenario 1: Only unlabeled data are available during training.
- Application scenario 2: Labeled training data are available from manual gating.

**Fig 2.**
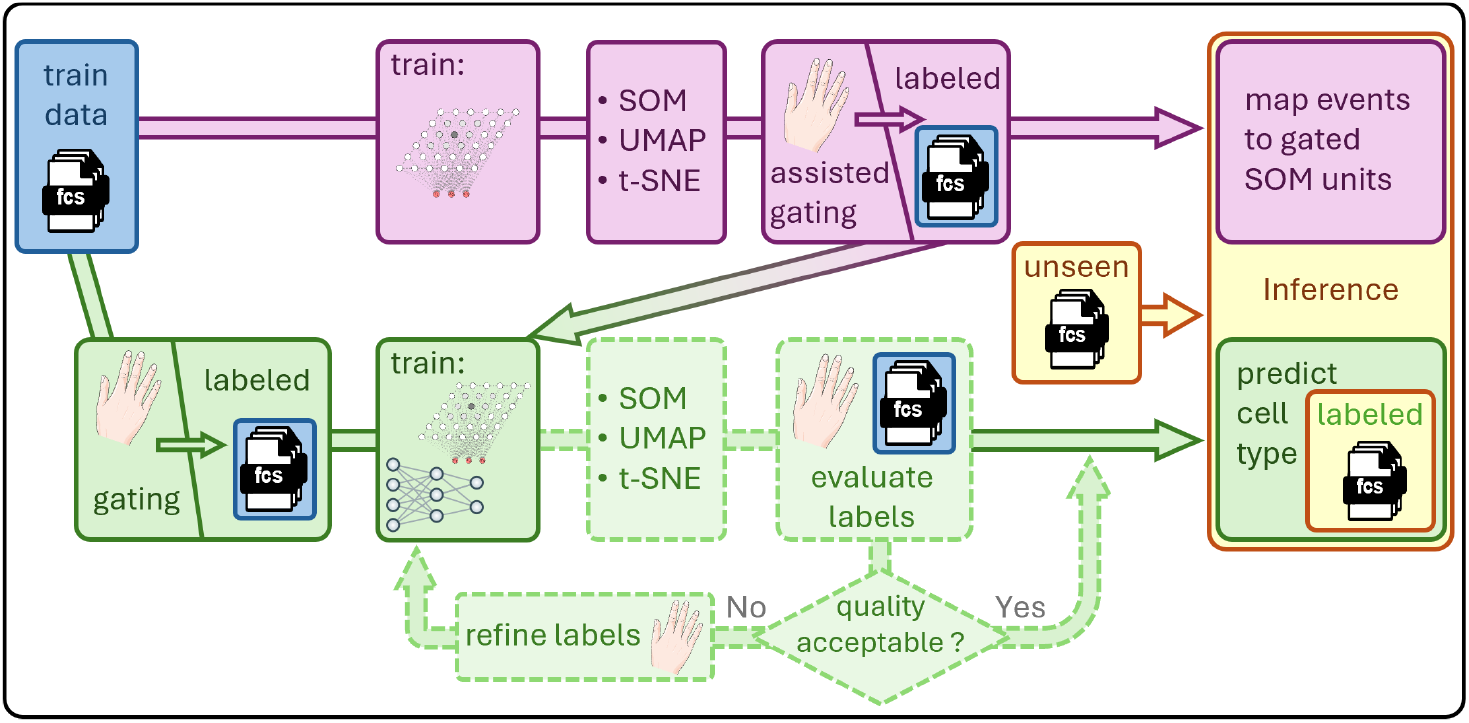
Overview of the FLAG-X workflow for gating clinical FCM data. Purple track: Unlabeled training data serves as the input for SOM training. Labeling is performed on the trained SOM and can be either used for re-mapping of new events or for crossover to the labeled data track. Green track: Labeled training data is provided by manual gating. An optional refinement loop for the ground truth labels is shown with dashed outline.

When no labeled training data are available (Fig 2, purple track), only the SOM component of the SOM classifier is trained, which does not require labeled training data. Each cell in the training data is mapped to its best matching unit in the SOM grid, which corresponds to a 2D grid coordinate. Additional 2D representations of the training data can be computed using PCA, UMAP [28], t-SNE [8]. The training data are annotated with these 2D representations and exported to an FCS file. Using conventional FCM software, practitioners can then assign cell labels by combining conventional FCM gating for selected training samples with ML-supported gating for all training samples in the 2D SOM, UMAP, or t-SNE representation spaces. The population labels thus defined can then either be used as ground truth for supervised model training (application scenario 2 visualized by crossover to the green track in Fig 2, see details below) or to annotate units of the trained SOM, enabling deduced labeling of new data through assignment of cells to best-matching SOM units.

If labeled training data are available (Fig 2, green track), the MLP and/or the SOM classifier are trained on the labeled samples. Optionally, 2D data representations (such as PCA, UMAP, t-SNE, and the mapping of events to units in the SOM grid) are added to the training data table. The augmented data table can then be exported to an FCS file, allowing the expert user to evaluate whether the population assignment of the events in the training data is plausible. If not, the labels can be refined and the training procedure repeated, thereby reinitiating the workflow with revised labeled data. During inference, the trained model assigns cell type labels to individual cells and again exports an FCS file, indicating the population labels and optional 2D representations. This output enables practitioners to easily visualize automated gating results with classical FCM analysis software tools (e. g., color-coding), recognize potential errors, and seamlessly continue with (manual) downstream analyses.

### FLAG-X implementations of the MLP and the SOM classifier are robust to limited training data

Fig 3 shows the results of our training data subsampling experiments. Both models implemented in FLAG-X maintained high performance even with limited training data, as test scores quickly approach the performance scores obtained with the full training set. With as few as 10–20 training samples, both models reached maximum performance. Beyond this point, performance fluctuated around the full-dataset results, with no consistent improvement. Additionally, the variability across test samples remained stable if few training samples were used, as shown by the empirical 95% confidence intervals plotted around each curve. Moreover, when manually curating the training subsets such that samples from each relevant phenotype are adequately represented, stable performance was achieved with fewer data compared to using randomly selected samples (Fig S6 in S1 Appendix).

**Fig 3.**
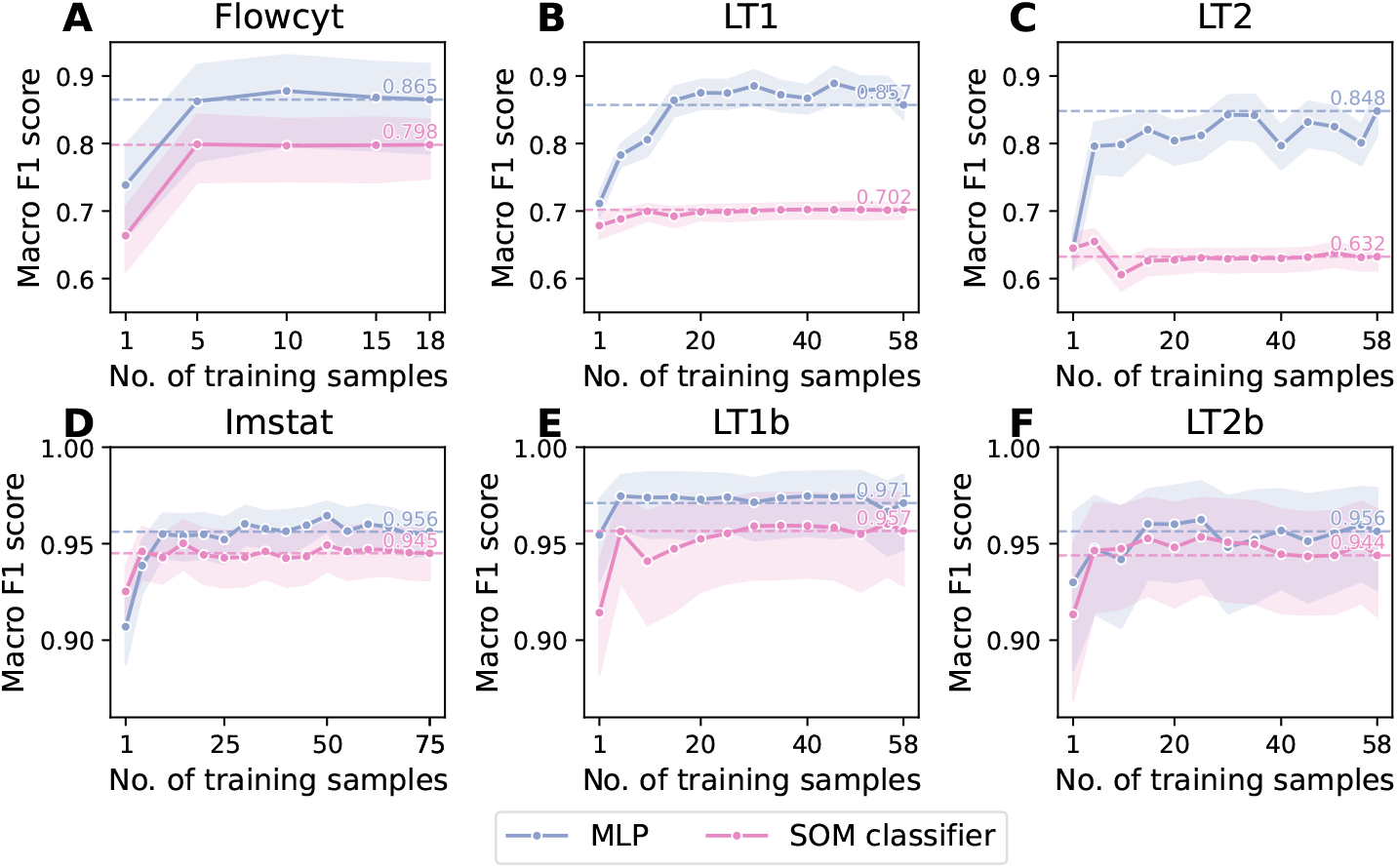
Average macro F1 scores across test samples of the MLP and the SOM classifier as a function of the number of training samples. Empirical 95 % confidence intervals for the sample-wise scores are shown around each curve. Performance achieved when training on the full dataset is visualized by horizontal lines. Y-axis range: 0.6–0.9 for A–C, 0.9–1.0 for D–F.

To establish a lower bound on the amount of training data required to achieve performance comparable to the full-dataset results, we trained the MLP and the SOM classifier on datasets where also the number of events per sample were systematically reduced. From these experiments, we determined the minimal number of samples and events for which the SOM classifier and the MLP achieve performances within the empirical 95% confidence interval of the full-dataset baseline by first minimizing over the number of samples (excluding the single-sample case) and subsequently over the number of events per sample. Although the MLP achieved superior performance when trained on the full dataset (see Fig 1 above), the SOM classifier reached its performance plateau with fewer data, particularly in datasets containing small cell populations such as Flowcyt, LT1, and LT2. The results are summarized in Table 1 and shown in full in Fig S7 in S1 Appendix.

**Table 1.** Minimal data requirements to match full-dataset performance.

| Datset | Flowcyt | Imstat | LT1 | LT2 | LT1b | LT2b |
| --- | --- | --- | --- | --- | --- | --- |
| MLP |  |  |  |  |  |  |
| No. of samples | 5 | 10 | 15 | 15 | 5 | 5 |
| No. of events per sample | 50,000 | 20,000 | all | all | 5000 | 5000 |
| Total no. of events | 250,000 | 200,000 | 1,414,575 | 1,434,948 | 25,000 | 25,000 |
| Minority count | 459 | 1864 | 5808 | 12,197 | 1801 | 1660 |
| SOM classifier |  |  |  |  |  |  |
| No. of samples | 5 | 5 | 5 | 5 | 5 | 5 |
| No. of events per sample | 20,000 | 5000 | 5000 | 5000 | 5000 | 5000 |
| Total no. of events | 100,000 | 25,000 | 25,000 | 25,000 | 25,000 | 25,000 |
| Minority count | 184 | 285 | 87 | 276 | 1801 | 1660 |

Since correct classification of the smallest class might be a limiting factor, we analyzed this quantitatively and identified the total number of events of the smallest class in the concatenated training samples (“minority count”) as an informative predictor for classifier performance. Full training set-level performance was achieved with a median minority count across datasets of roughly 2000 for the MLP and below 1000 for the SOM classifier, showing that both methods and especially the SOM classifier are data-efficient models. For detailed results, we refer to Table 1 and Fig S8 in S1 Appendix.

### FLAG-X enables model training on local machines

Given that high performance can be achieved with few labeled samples of reasonable size, FLAG-X enables training on local machines equipped with commonly available CPUs and GPUs, which is critical for practical deployment in clinical settings. Models trained locally, with reduced number of training samples and per-sample downsampling as previously determined to be sufficient to match the full-dataset baseline (see Table 1), achieved performance comparable to those trained using high-performance computing resources on the full datasets. Also, training times and memory requirements remained low, below 20 minutes and 10 Gigabyte (see Table 2), further supporting the feasibility of local deployment. Detailed performance scores are provided in Fig S9 in S1 Appendix.

**Table 2.** Wall time and peak memory usage during model training and inference with a standard local machine. Training the MLP classifier required 0.12 GB of peak GPU memory during training across all datasets.

|  | Flowcyt | Imstat | LT1 | LT2 | LT1b | LT2b |
| --- | --- | --- | --- | --- | --- | --- |
| Training – Wall Time |  |  |  |  |  |  |
| MLP | 4 m 30 s | 3 m 50 s | 14 m 53 s | 16 m 21 s | 19 s | 15 s |
| SOM clf | 3 m 40 s | 56 s | 56 s | 58 s | 57 s | 58 s |
| Training – Peak Memory (CPU) |  |  |  |  |  |  |
| MLP | 6.30 GB | 10.37 GB | 9.26 GB | 10.16 GB | 9.76 GB | 8.92 GB |
| SOM clf | 1.13 GB | 1.44 GB | 1.05 GB | 1.16 GB | 1.21 GB | 1.21 GB |
| Inference – Wall Time, (mean ± std) | | | | | | |
| MLP | 0.63 ± 0.20 s | 0.03 ± 0.05 s | 0.05 ± 0.05 s | 0.06 ± 0.06 s | 0.05 ± 0.05 s | 0.05 ± 0.05 s |
| SOM clf | 5.70 ± 1.74 s | 0.28 ± 0.12 s | 0.72 ± 0.09 s | 0.72 ± 0.11 s | 0.72 ± 0.09 s | 0.71 ± 0.09 s |
| Inference – Peak Memory (CPU), (mean ± std) | | | | | | |
| MLP | 2.49 ± 0.22 GB | 1.52 ± 0.01 GB | 1.79 ± 0.01 GB | 1.93 ± 0.02 GB | 1.65 ± 0.02 GB | 1.67 ± 0.02 GB |
| SOM clf | 3.85 ± 0.77 GB | 1.13 ± 0.08 GB | 1.40 ± 0.04 GB | 1.52 ± 0.05 GB | 1.52 ± 0.05 GB | 1.52 ± 0.05 GB |

### FLAG-X achieves favorable recall-precision trade-offs in clinical subpopulation selection tasks

In clinical workflows, downstream analyses often focus on a specific cell population, such as B cells in the context of lymphoma classification or immature myeloid cells in the case of measurable residual disease determination in AML. In this binary setting, prioritizing high recall during preliminary gating for the cell population of interest ensures that clinically relevant cells are not missed. To evaluate FLAG-X in such a scenario, we analyzed the binary recall and precision of the locally trained models on the LT1b and LT2b datasets, i. e., “B cell” versus “all other cells” classification. Both the MLP and the SOM classifier exhibited favorable recall-precision trade-offs, achieving recall scores *≥*0.85 at threshold levels where precision remained higher than recall (Fig 4). These results demonstrate that FLAG-X can be effectively applied in clinically relevant, recall-oriented use cases without requiring large annotated datasets.

**Fig 4.**
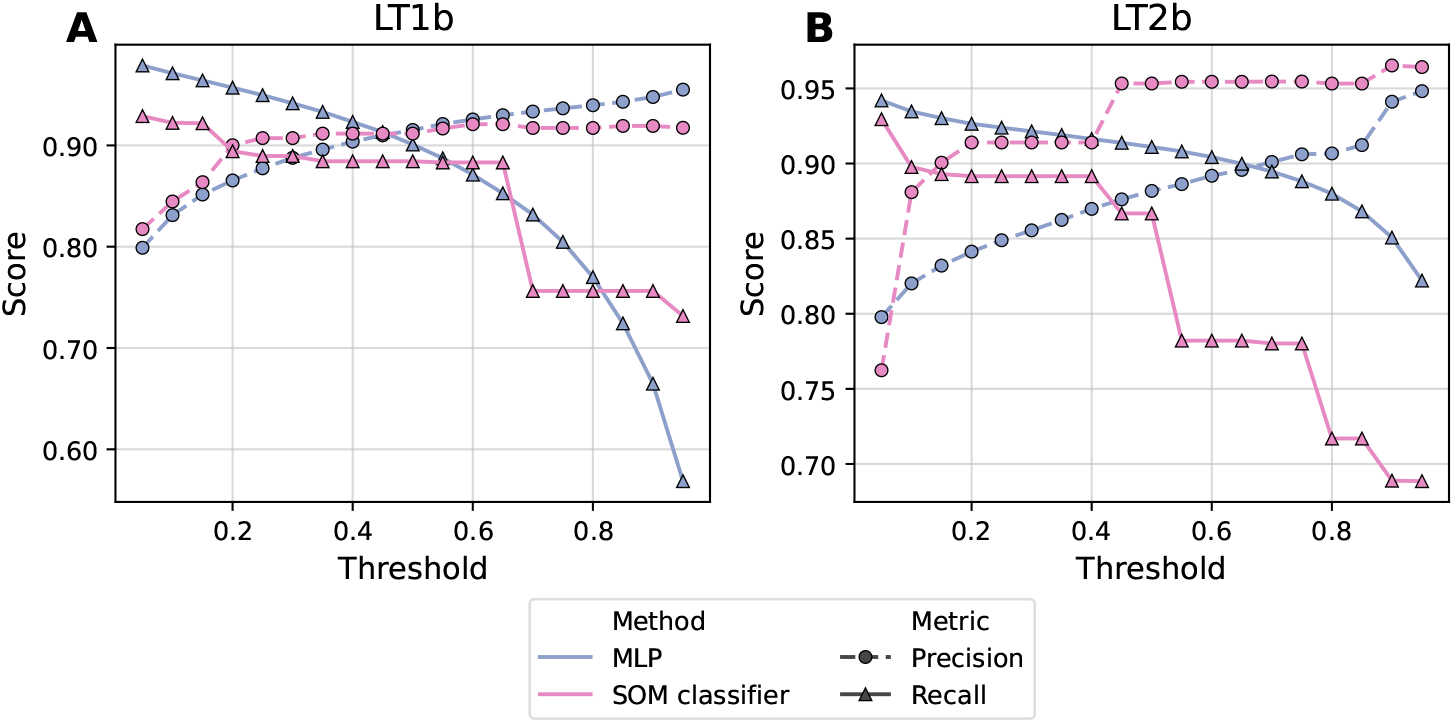
Binary recall and precision scores at varying probability thresholds, averaged across test samples. Results for the LT1b dataset (A) and for the LT2b dataset (B). Shown probability thresholds are 0.05, 0.1, …, 0.9, 0.95. Method is indicated by color, performance metric is indicated by marker and line type.

### Automated gating with FLAG-X remains robust for target populations with diverse phenotypes

Next, we investigated how automated gating with FLAG-X performs when the target cell population is phenotypically diverse. For this, we trained SOM and MLP models on a subset of the LT1b dataset comprising samples from different lymphoma subtypes and from the normal controls. While the ground truth B cell populations were defined as “CD45+ and CD19+/(+) and CD79b/CD4+/(+)” with manual adjustment of gates for each training sample, B cells originating from training samples belonging to different lymphoma subtypes grouped into distinct clusters in the UMAP and SOM representations of the concatenated training data (Fig 5A, B). Particularly, the trained SOM preserved heterogeneity across lymphoma subtypes alongside the ground truth cell populations, as different lymphoma subtypes were represented by distinct units of the trained SOM. This shows that automated gating with the SOM classifier can preserve diversity of the target population, and that linking SOM units to biological phenotypes at the sample level may even provide a basis for subsequent downstream analysis (subclassification of B-cell lymphomas in this example).

**Fig 5.**
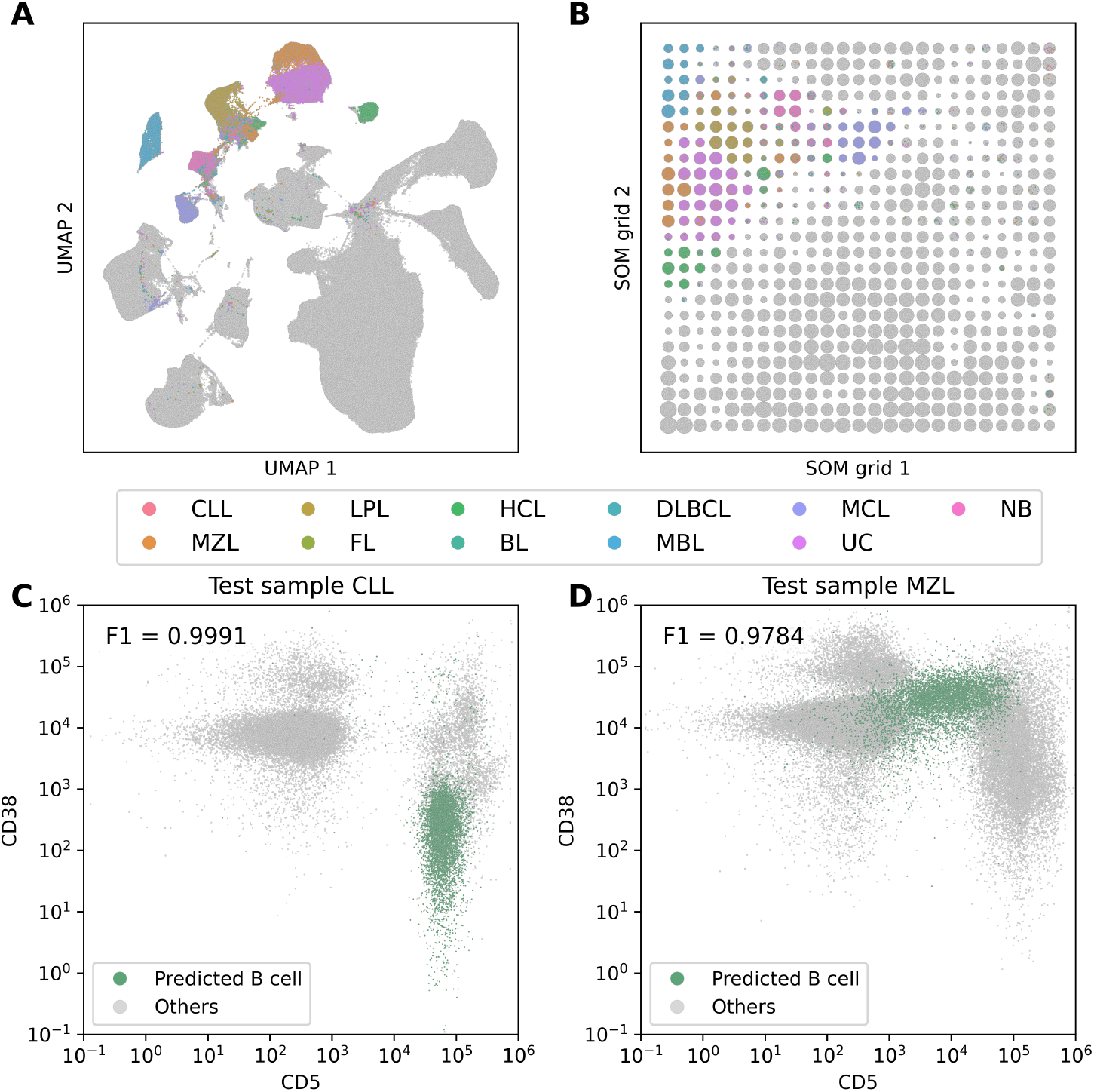
Automated gating of B cells across diverse lymphoma subtypes. (A, B) UMAP and SOM representations of training samples. Colored dots represent B cells (ground truth cell type labels) and color indicates lymphoma subtype. (C, D) Scatter plots of CD5 vs. CD38 expression of exemplary CLL and MZL test samples. B cell predicted by the MLP are highlighted in green. The F1 scores (binary, positive class = B cells) quantify sample-specific gating performance of the trained MLP.

Unlike the SOM classifier, the MLP does not provide an intrinsic representation of population structure. To assess whether automated gating results produced by the MLP also remain meaningful when the target population is diverse, we applied the trained MLP to gate samples from a reduced test set containing one sample of each lymphoma subtype. Heterogeneity across test samples is exemplified in Fig 5C and D by a scatter plot of CD5 vs. CD38 expression, for a chronic lymphocytic leukemia (CLL) and marginal zone lymphoma (MZL) test sample with predicted B cells highlighted in green (corresponding plots for test samples from the other lymphoma subtypes are shown in Fig S5 in S1 Appendix). The automatically gated B cells from the CLL and MZL samples exhibit very different phenotypes. For instance, CD5 expression is higher in the CLL sample than in the MZL sample, consistent with existing literature [29]. This demonstrates that the MLP can correctly identify cells of diverse phenotypes when the target population is heterogeneous.

### Usage example: Iterative ground truth label refinement with FLAG-X

Finally, we illustrate the iterative improvement of ground truth training labels on the Imstat dataset, where NK cells were defined as “(CD16+CD45++ or CD56+CD45++) and not (Monocytes or Granulocytes or CD3+)” by classical gating. In the trained SOM as well as in the UMAP representation of the training data, the core NK population is clearly separated from other events, while outliers can be recognized as likely artifacts of imprecise ground truth gating (marked with red arrows in Fig 6A, B). Refining the ground truth gating to the core NK population in the trained SOM (Fig 6C) yields a clear improvement, as this adjustment also substantially reduces the number of outlier NK cells in the UMAP representation (Fig 6D).

**Fig 6.**
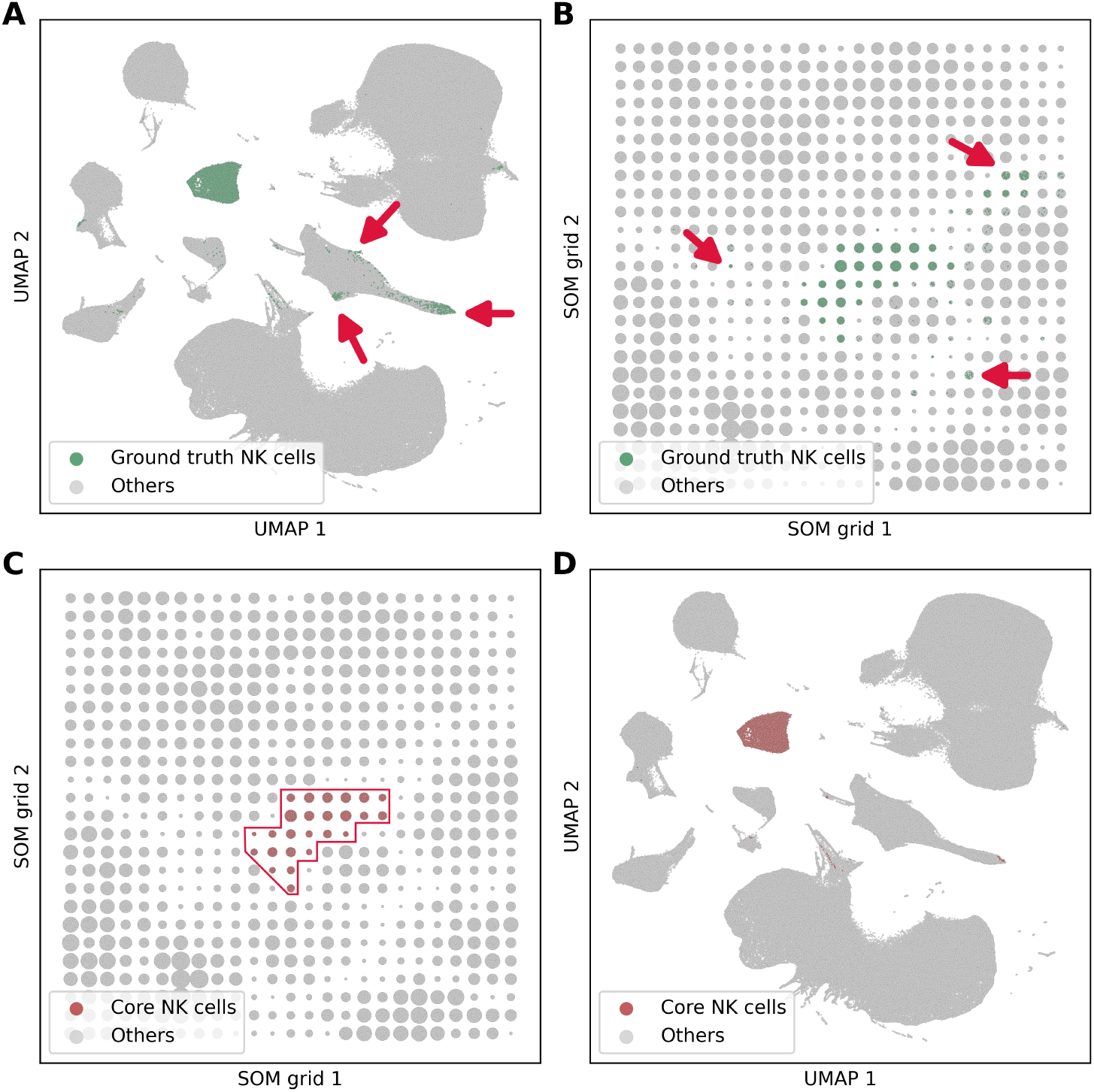
Illustration of iterative refinement of “ground truth” population assignment for NK cells in the Imstat dataset. (A, B) UMAP and SOM representation of the training data with “ground truth” NK cells shown in green. Outliers outside of the core NK population are highlighted with red arrows. (C) Refined ground truth gating defined as the core NK population in the trained SOM (marked by a red polygon). (D) Refined ground truth gating shown in UMAP representation.

## Discussion

Our comparative evaluation of supervised gating methods for FCM data revealed several key findings. Most importantly, the two simplest models, the SOM and MLP classifiers, performed very well and reached clinically useful gating performances on unseen data (Fig 1). Additional architectural complexity such as DGCyTOF’s correlation-based calibration of MLP predictions did not lead to improved performance but substantially increased resource requirements. In view of these results, we asked if the two simple models can reach strong predictive performance even when trained on annotated data from only few samples, possibly sub-sampled to few cells per sample. Indeed, our analyses yielded positive results: For all combination of models and test datasets, a maximum of 15 training samples was sufficient to reach performances obtained with the full training sets, often with as few as 5000 cells per sample, as long as the smallest class of interest is represented by a sufficient number of cells (Table 1). We then showed that, when using such subsampled training data, both the MLP and the SOM classifier can be trained on local machines within minutes, without compromising predictive performance.

Taken together, these empirical findings show that clinical deployment of ML-based gating methods for FCM data is in reach. To further advance towards translation, we developed the FLAG-X Python package. FLAG-X not only provides a uniform and user-friendly interface for the SOM and MLP classifiers, but also supports hybrid expert-in-the-loop workflows, input and export of FCS files for seamless integration with standard FCM data analysis software, as well as subsampling methods to enable local and resource-efficient model training. Starting with as few as 5–20 samples for training, new samples can be annotated in very little time, enabling fast analysis of large collections of datasets.

Our study opens up several interesting directions for future work. First of all, the cell type predictions of the SOM classifier and the MLP on unseen test data aligned very well but not perfectly with cell type labels obtained via manual gating. It is possible that the predictive performances of the MLP and (closely behind) SOM models have already reached the upper bounds imposed by the uncertainty in the manual training data annotation, especially in view of the high inter-rater variability in FCM gating observed in previous studies [3]. In this context, our proposed expert-in-the-loop workflow, integrating 2D representations for gating, can improve classification of the training data, and consequently the correct classification of new data. Alternatively, a further improvement in predictive performance could be achieved through more complex models, e. g., using self-supervised pre-training as in recent transformer-based models that support cell type prediction for single-cell RNA-sequencing data [30, 31]. This question would have to be tested in a follow-up study, ideally with several expert annotators who independently gate the same data and then qualitatively compare ML-based cell type predictions to manual cell type labels provided by the other annotators.

## Conclusion

This study shows that prudent combination of comparatively simple methods is sufficient to perform reliable population annotation of FCM data while requiring only moderate amounts of training data and computational resources, thereby providing a solid basis for clinical translation. To further advance towards automated gating, we propose hybrid workflows, implemented in the FLAG-X Python package, that facilitate the integration of ML methods into existing clinical workflows. In general, further efforts to automate gating as well as subsequent downstream analyses should not focus solely on method development, but should also consider the practical requirements for clinical adoption.

## Supporting information

**S1 Appendix. Supplementary information**.

## Author contributions

**Conceptualization:** Paul Martini, David B. Blumenthal, Stefan W. Krause

**Data Curation:** Marziyeh Mohammadi, Michael C. Thrun, Stefan W. Krause

**Formal Analysis:** Paul Martini

**Funding Acquisition:** David B. Blumenthal, Stefan W. Krause

**Investigation:** Paul Martini

**Methodology:** Paul Martini, David B. Blumenthal, Stefan W. Krause

**Software:** Paul Martini, Marziyeh Mohammadi

**Supervision:** David B. Blumenthal, Stefan W. Krause

**Validation:** Paul Martini

**Visualization:** Paul Martini

**Writing – Original Draft Preparation:** Paul Martini

**Writing – Review & Editing:** Paul Martini, Marziyeh Mohammadi, Michael C. Thrun, David B. Blumenthal, Stefan W. Krause

## Availability statement

- The Flowcyt dataset is available at https://cuicloud.unige.ch/index.php/s/55PHBLEynrp5pN8. Instructions and code for data processing are available at https://github.com/LorenzoBini4/FlowCyt-Classification-Benchmark.
- The Imstat, LT1, and LT2 datasets are not publicly available due to data protection regulations and ethical restrictions. Access to these datasets can only be granted upon dedicated applications to the Ethics Commissions of, respectively, the Faculty of Medicine of the Friedrich-Alexander-Universität Erlangen-Nürnberg (for the Imstat dataset) and the School of Medicine of the Philipps-Universität Marburg (for the LT1 and LT2 datasets). For information on how to initiate this process, please contact the corresponding author S. W. K.
- The FLAG-X Python package is available at https://anaconda.org/channels/bioconda/packages/flagx/overview and https://github.com/bionetslab/FLAG-X. The GitHub repository also contains the Jupyter notebooks that exemplify FLAG-X’s unsupervised and supervised workflows.
- Source code to reproduce the results reported in this article is available at https://github.com/bionetslab/FLAG-X-validation.

## Ethics statement

Data from routine measurements were analyzed in accordance with legal regulations which permit the internal use of pre-existing data for research and analytical purposes.

## Acknowledgments

The authors gratefully acknowledge the scientific support and HPC resources provided by the Erlangen National High Performance Computing Center (NHR@FAU) of the Friedrich-Alexander-Universität Erlangen-Nürnberg (FAU). The hardware is funded by the German Research Foundation (DFG).

