## Supplementary material for "FLAG-X: Hybrid machine learning workflows for automated gating of clinical flow cytometry data": S1_Appendix-Supplementary_information

#### S1 Self-organizing maps and SOM classifier

**Self-organizing maps.** A self-organizing map (SOM) consists of units $u = 1,\dots,U$ which are arranged in a two-dimensional plane, typically in a rectangular or hexagonal grid. Each unit $u$ is characterized by

- a fixed position in the grid, represented as a two-dimensional coordinate $\text{pos}(u) = (i(u),\, j(u))$,
- and a learnable weight vector $\mathbf{u} \in \mathbb{R}^m$.

The dimension $m$ of the SOM's weight vectors matches the number of features $m'$ of the vectorial input data $X \in \mathbb{R}^{n\times m'}$. The standard SOM training procedure is outlined in Algorithm S1. Before training, the SOM's weight vectors are initialized randomly. For each training sample $X_{j\bullet}$ its best matching unit (BMU), meaning the unit whose weight vector $\mathbf{u}$ is most similar to the sample, is identified. The BMU, along with units in its grid neighborhood, are then updated to more closely resemble the current sample. Typically, the magnitude of the update decreases with increasing distance from the BMU and is further controlled by the learning rate function $\alpha$. These iterative updates are performed for $T$ passes over the training set (epochs). The neighborhood function $h$ also depends on the radius function $r$. Both, the learning rate and the radius function usually decrease with the number of completed epochs. Commonly used neighborhood functions are

- bubble: $h(\text{pos}(u), \text{pos}(u'), r) = \begin{cases} 1 & \text{if } \|\text{pos}(u) - \text{pos}(u')\|_2 \le r \\ 0 & \text{otherwise,} \end{cases}$,
- and Gaussian: $h(\text{pos}(u), \text{pos}(u'), r) = \exp(\frac{-\|\text{pos}(u)-\text{pos}(u')\|_2^2}{2(\sigma r)^2})$.

SOMs learn in an unsupervised, competitive fashion such that the trained SOM represents a lower dimensional map of the input space where similar samples are mapped closer together and units that are BMU for comparatively many samples can be associated with dense regions in the input space.

**SOM classifier.** Given labeled input data $(X,y) \in \mathbb{R}^{n\times m} \times \{1,\dots,l\}^n$, $l \in \mathbb{N}$, SOMs can be extended for supervised classification. This is achieved by analyzing the label composition of SOM units after training. Let $y(u) \in \{1,\dots,l\}^{n(u)}$, $n(u) \le n$ be the label vector corresponding to samples of the training set that have SOM unit $u$ as BMU after SOM training. Then the class fractions at unit $u$ are

$$c_k(u) = \frac{1}{n(u)} \sum_{j=1}^{n(u)} \mathbb{1}(y(u)_j = k) \quad \text{for } k \in \{1,\dots,l\}.$$

The majority class $k^*(u) = \text{argmax}_{k\in\{1,\dots,l\}}(c_k(u))$ is then assigned to be the label of unit $u$. When presented with a previously unseen sample $x \in \mathbb{R}^m$ SOM classifier makes a prediction by assigning it the majority class $k^*(\text{bmu}(x))$ of its BMU. The class fractions at the best matching unit $c_k(\text{bmu}(x))$, $k \in \{1,\dots,l\}$ can be interpreted as as probabilistic prediction across all classes.

**Algorithm S1** SOM Training

**Input:** Data $X \in \mathbb{R}^{n \times m}$, SOM, learning rate function $\alpha : \mathbb{N} \to \mathbb{R}_{\geq 0}$, radius function $r : \mathbb{N} \to \mathbb{R}_{\geq 0}$, neighborhood function $h : \mathbb{R}^2 \times \mathbb{R}^2 \times \mathbb{R} \to \mathbb{R}_{\geq 0}$, number of epochs $T \in \mathbb{N}$
**Initialize** weight vectors randomly
**for** epochs $t = 1, \dots, T$ **do**
  **for** samples $X_{j\bullet}$, $j = 1, \dots, m$ **do**
    Find BMU: $\text{bmu}(X_{j\bullet}) = \text{argmin}_{u \in \{1,\dots,U\}}(\|X_{j\bullet} - \mathbf{u}\|_2)$
    **for** SOM unit $u = 1, \dots, U$ **do**
      Compute neighborhood: $h_{u,\text{bmu}(X_{j\bullet})} = h(\text{pos}(\text{bmu}(X_{j\bullet})), \text{pos}(u), r(t))$
      Update weight vector: $\mathbf{u} \leftarrow \mathbf{u} + \alpha(t) \cdot h_{u,\text{bmu}(X_{j\bullet})} \cdot (X_{j\bullet} - \mathbf{u})$
    **end for**
  **end for**
**end for**
**Output:** Trained SOM

### S2 Qualitative assessment: interpretability, user-friendliness, and practical feasibility

In addition to gating performance, we evaluated the four methods on qualitative criteria that are critical for practical deployment in clinical and research settings, leveraging our synergistic experience in ML model development and clinical FCM data analysis: interpretability, user-friendliness, and practical feasibility of deployment in a clinical setting. Table S1 gives a summary of our findings; more detailed explanations are provided below.

**Interpretability.** GateMeClass discretizes marker expression into categorical levels and uses rule-based decision tables, mimicking manual gating strategies. DGCyTOF builds on MLP with an additional feedback calibration step, but the internal decision process remains opaque for both methods. For the SOM classifier, the cell-type fractions at the best matching unit allow insight into the decision process. Further, this distribution of cell types across SOM units can be interpreted as a prediction confidence. The mapping of events to SOM units is a built-in 2D visualization. In contrast, the MLP provides probabilistic outputs that are not inherently interpretable and does not offer built-in visualization.

**User-friendliness.** GateMeClass is implemented in R and integrates well with R-based analysis workflows. DGCyTOF, the MLP, and the SOM classifier are implemented in Python. DGCyTOF and the MLP require installation of Pytorch-based deep learning frameworks. Our implementation of SOM classifier includes a command-line interface that can be used with minimal programming experience. All tools are publicly available as open-source repositories with documentation on GitHub.

**Practical feasibility.** When a gating decision table is provided by the user, GateMeClass does not require labeled training data. Both DGCyTOF and the MLP cannot be trained without labeled training data. The SOM component of the SOM classifier is trained in an unsupervised fashion; labels are required only for the assignment of cell types to SOM units. During evaluation on real-world datasets, we encountered issues with GateMeClass and DGCyTOF:

- GateMeClass failed when applied to data with lower-bound transformations, such as $f(x) = \log_{10}(x)$ if $x > 100$, $\log_{10}(100)$ otherwise. Failure occurred during gaussian mixture model fitting, likely due to constant values in the distribution tail introduced by the cutoff.
- DGCyTOF produced out-of-memory errors during inference on FCM datasets of typical size, due to the high memory cost of computing large correlation matrices (for details see Section S2.1).

**Table S1.** Qualitative assessment of GateMeClass, DGCyTOF, the MLP, and the SOM classifier. Symbols: $+$ (positive), $o$ (neutral), $-$ (negative).

| Category | Subcategory | GateMeClass | DGCyTOF | MLP | SOM classifier |
|---|---|---|---|---|---|
| Interpretability | Human-readable decisions | $+$ | $-$ | $-$ | $o$ |
| | Prediction confidence provided | $-$ | $-$ | $o$ | $+$ |
| | Integrated visualization | $-$ | $-$ | $-$ | $+$ |
| User-friendliness | Programming language | R | Python | Python | Python |
| | Setup complexity | $+$ | $-$ | $o$ | $+$ |
| Practical feasibility | Requires labeled training data | depends | yes | yes | depends |
| | Robustness to data transformations | $-$ | $+$ | $+$ | $+$ |
| | Handling of real-world dataset sizes | $+$ | $-$ | $+$ | $+$ |
| | Training time | $+$ | $+$ | $+$ | $+$ |
| | Inference time | $-$ | $-$ | $+$ | $+$ |

Training times were under 3 hours for all methods with training datasets of up to 11,671,354 cells using high-performance computing resources. The MLP and the SOM classifier had average per-sample inference times typically below one second and 30 seconds respectively, while DGCyTOF and GateMeClass required up to 10 minutes. Detailed training and inference times are reported in Table S10, and details on the computational environment are provided in the Methods section of the main manuscript.

#### S2.1 DGCyTOF out of memory error

At inference time Spearman correlation matrices are computed at two points:

(1) For each cell type $c \in \{c_1, \dots, c_k\}$ present in the training data the pairwise correlations are computed among cells predicted with high probability to belong to $c$. The resulting correlation matrix has a memory requirement of $O(\max_k\{|\{\text{cells belonging to } c_k\}|\}^2)$.

(2) For cells classified with low confidence, denoted as $C_{\text{low}} = \{x_1, \dots, x_q\}$, Spearman correlations are computed between each such cell and the high-confidence cells of each cell type. This has memory requirement $O(|C_{\text{low}}| \cdot |\{\text{cells belonging to } c_k\}|)$.

A single flow cytometry sample typically contains up to 100,000 individual events, with the majority class accounting for as much as 80% of the total population (see Figures S1, S2). Consequently, the correlation matrix computed at (1) is an $80{,}000 \times 80{,}000$ matrix. At 64 bit floating point precision, this requires $80{,}000^2 \cdot 8\,\text{Byte} = 51.2\,\text{GB}$ of memory, which exceeds the capacity of most standard computing environments. To address this, the authors of DGCyTOF implemented a hard cutoff of 12,000 cells for computing correlation matrices for a specific class (label 1), which appears to correspond to the majority class in their datasets (compare: `https://`

github.com/lijcheng12/DGCyTOF/blob/main/DGCyTOF_Package/DGCyTOF/__init__.py, lines 228–232, accessed 14.05.2025). To avoid out of memory errors when applying DGCyTOF to our datasets, we extended this cutoff to all classes.

### S3 Supplementary Figures

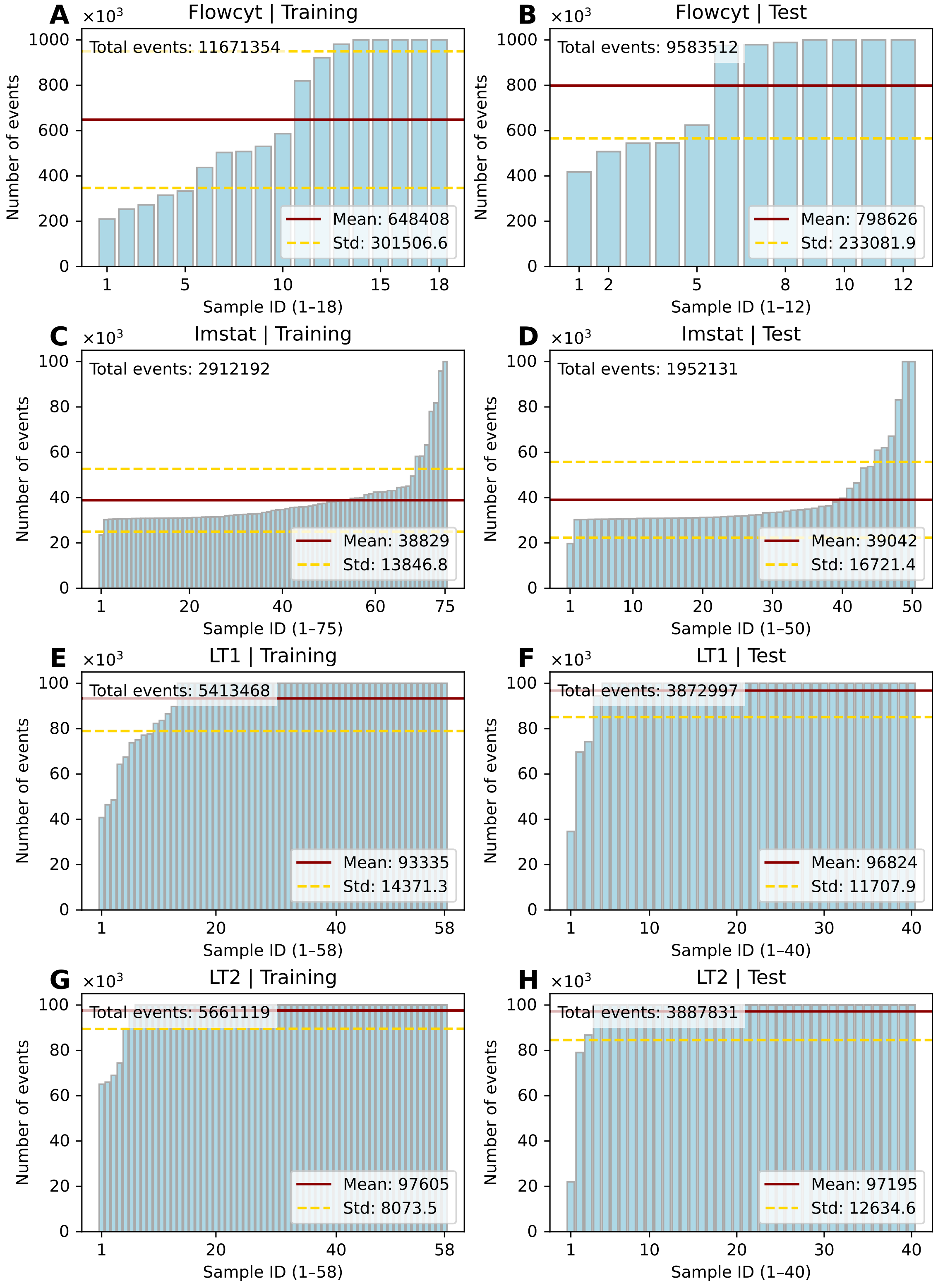


**Fig S1.** Bar plots of sample sizes in the training and test sets.

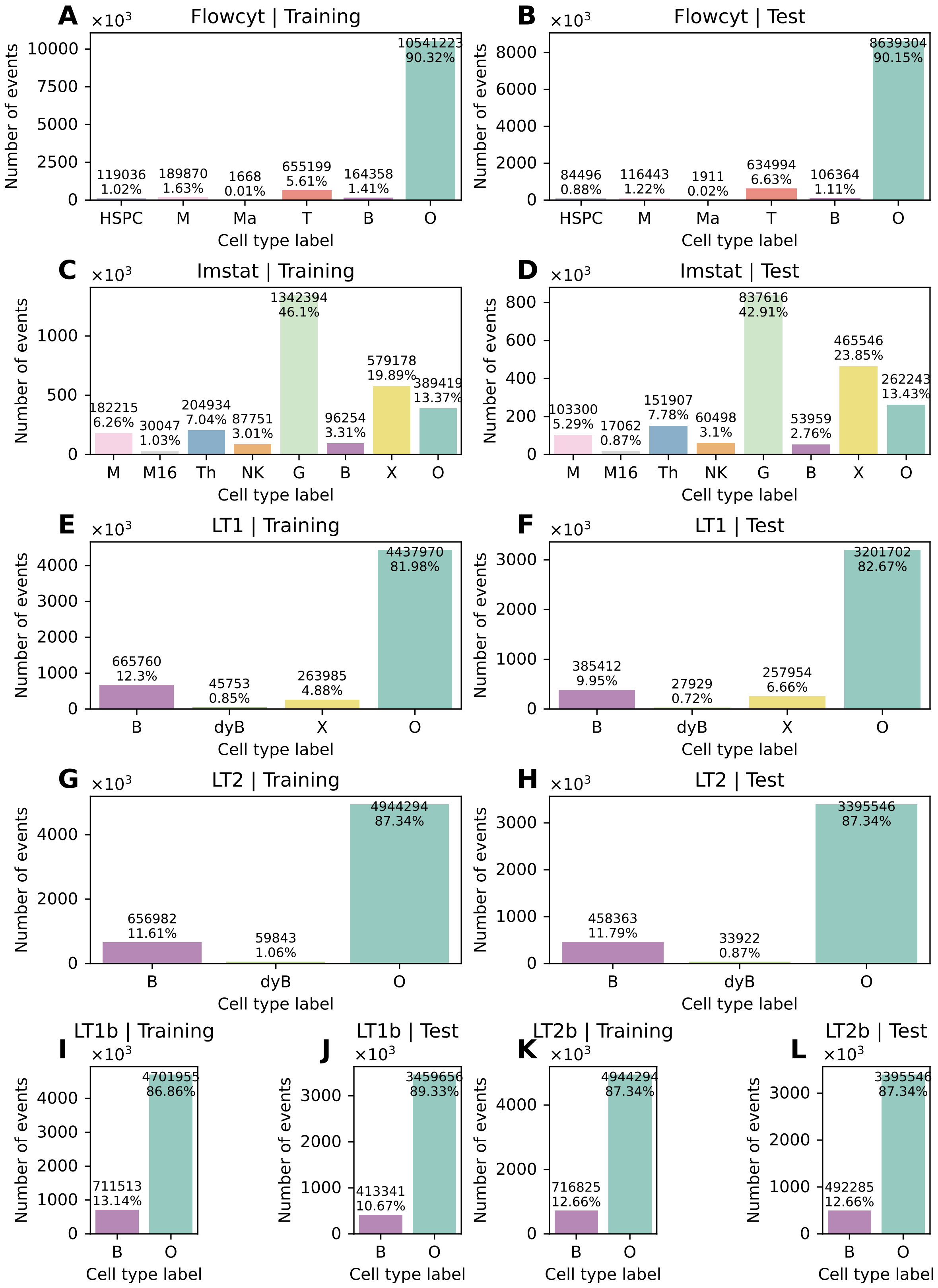


**Fig S2.** Class balances in the training and test sets.

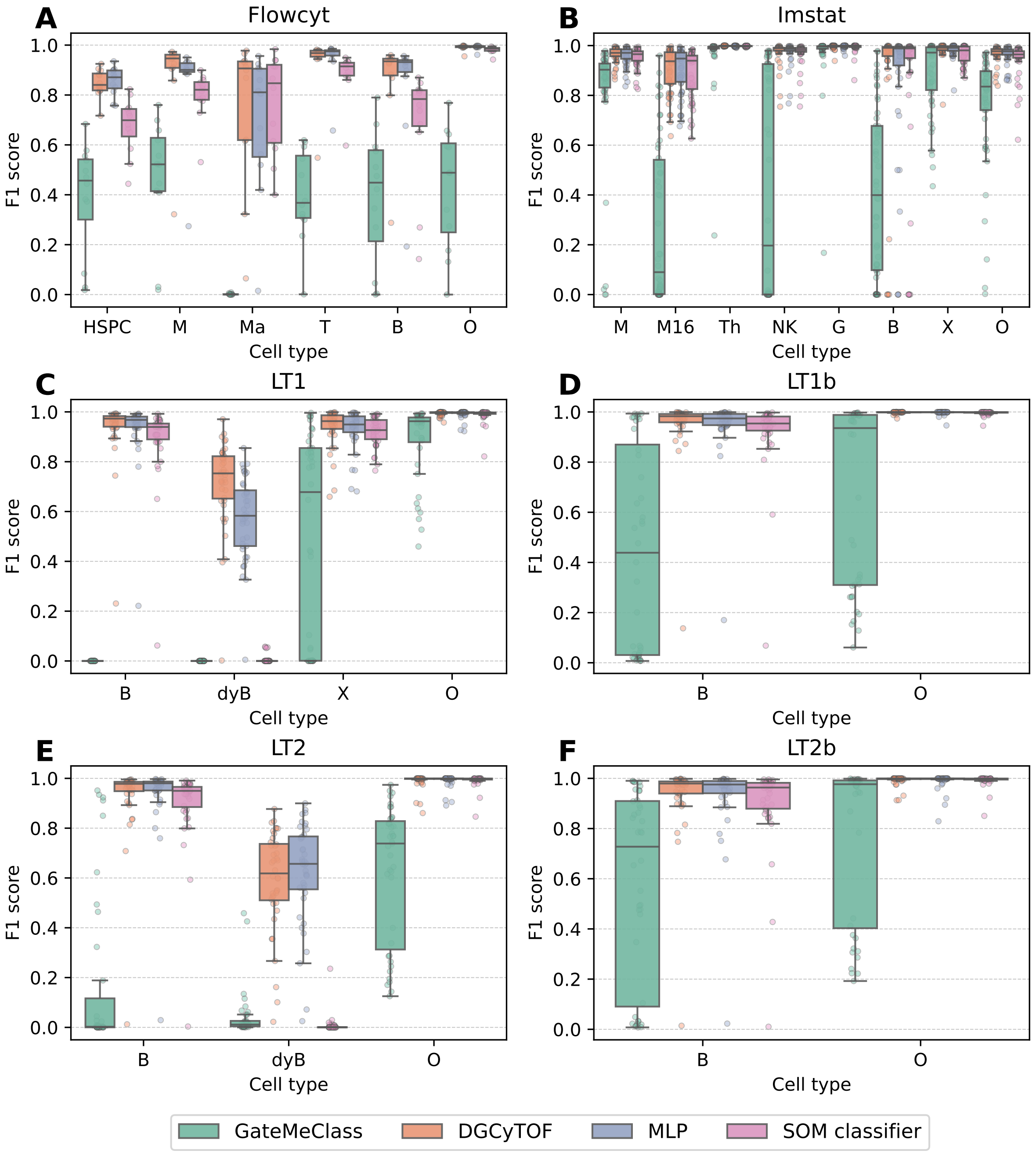


**Fig S3.** Class-wise F1 scores, reported for individual test samples across datasets. GateMeClass, and to a lesser extent SOM classifier, consistently underperform on minority classes, e.g., M16, B, and NK in the Imstat dataset (A), and classes Ma, HSPC, B, and M in the Flowcyt dataset (B). In the Lymphoma datasets performance increases markedly when subpopulations are merged with their parent populations, as seen by comparing (C) to (D) and (E) to (F).

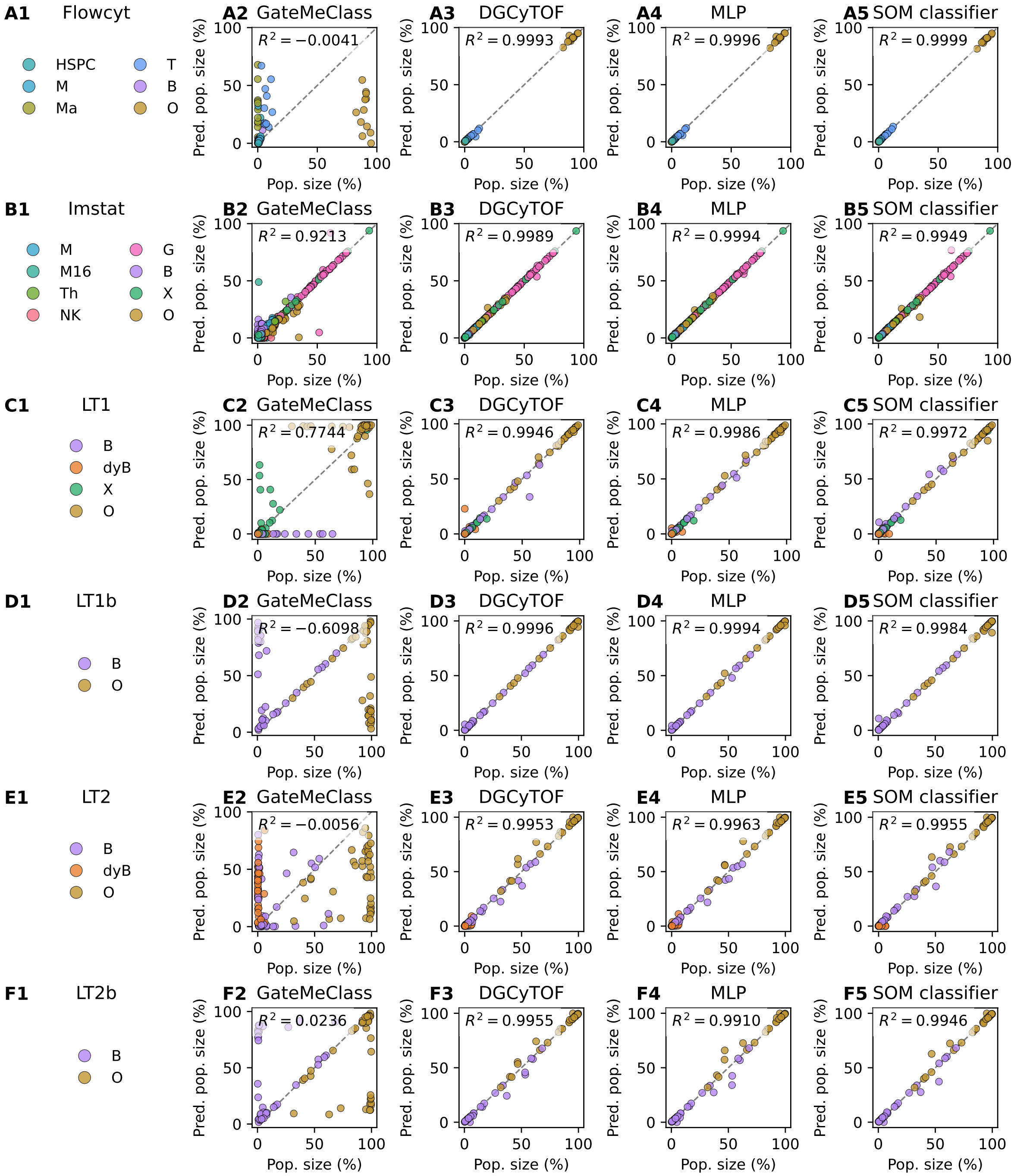


**Fig S4.** Predicted vs. true cell population sizes for the LT1, LT1b, LT2, LT2b, and Flowcyt datasets. Each point corresponds to one cell type from one sample. The coefficient of determination ($R^2 \in (-\infty, 1]$, higher is better) with respect to the identity line was used to assess goodness-of-fit. All methods except GateMeClass show near-perfect alignment with the ground truth, with $R^2$ exceeding 0.99, while GateMeClass trails behind with a maximal $R^2$ of 0.9213 across datasets.

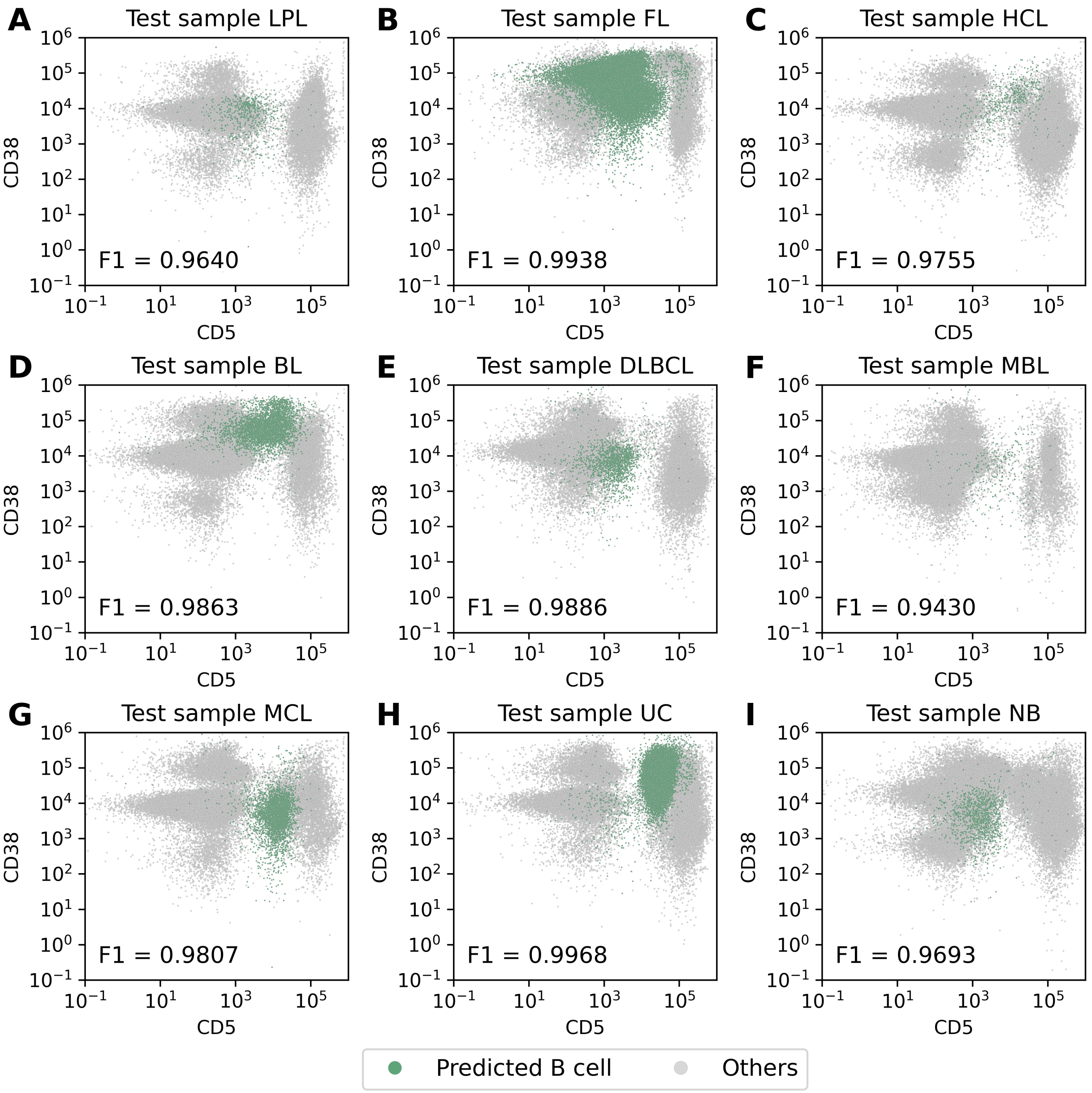


**Fig S5.** Scatter plots of CD5 vs. CD38 expression of test samples from diverse lymphoma subtypes. B cell predicted by the MLP are highlighted in green. The F1 scores (binary, positive class = B cells) quantify sample-specific gating performance of the trained MLP.

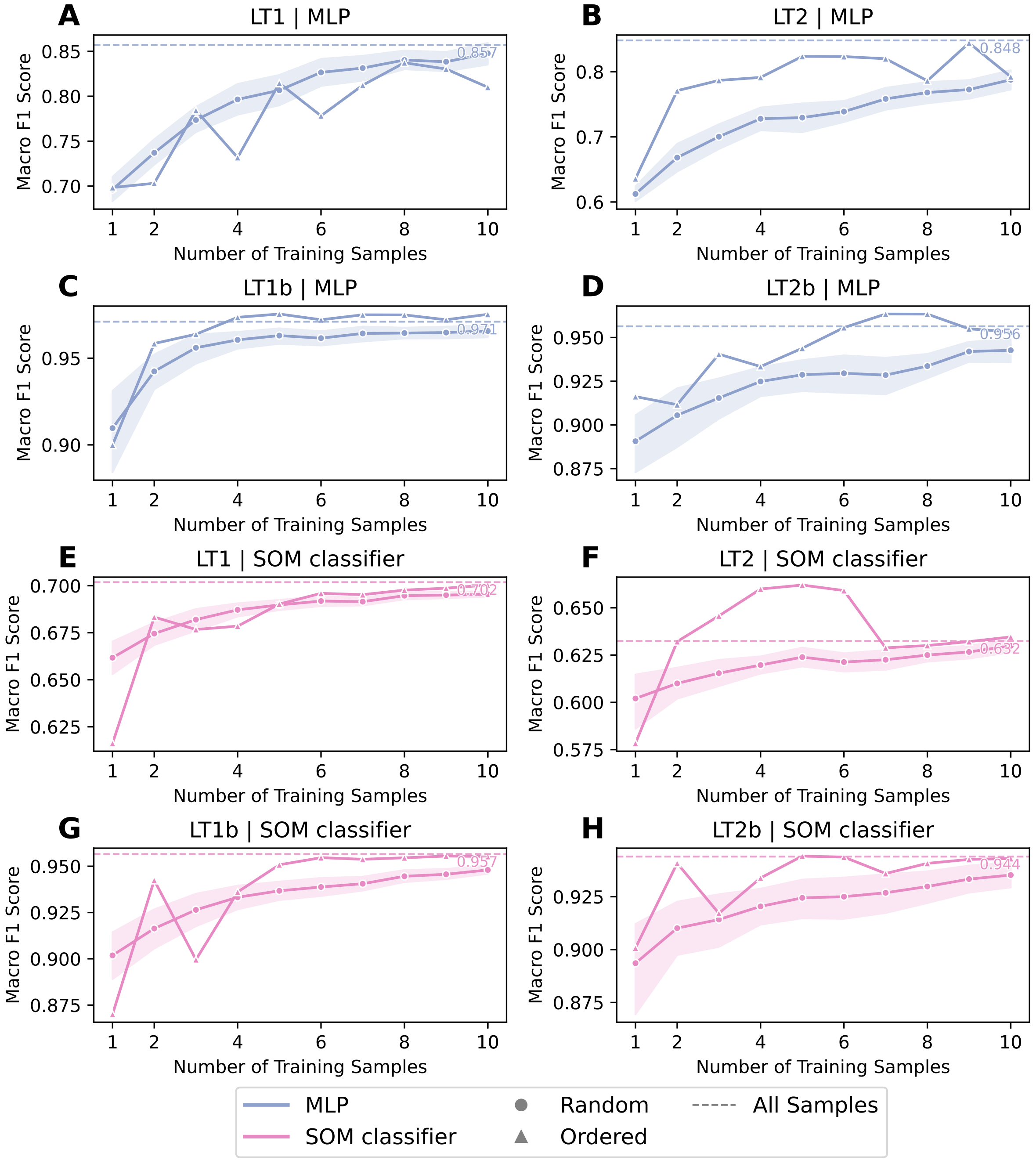


**Fig S6.** Average macro F1 scores across test samples of MLP and SOM classifier for varying numbers of training samples. Triangle markers show results for a predefined sample order, dots represent the mean performance across $n = 30$ trials with random sample orders. Shaded areas indicate empirical 95% confidence intervals for the scores across trials. Performance with the full training set is shown as a dashed line. With the exception of the LT1 dataset, convergence towards the all-sample performance is faster with a predefined sample order. This suggests that, when resources for labeling samples are limited, curating a training set in which diverse phenotypes are represented can improve model performance.

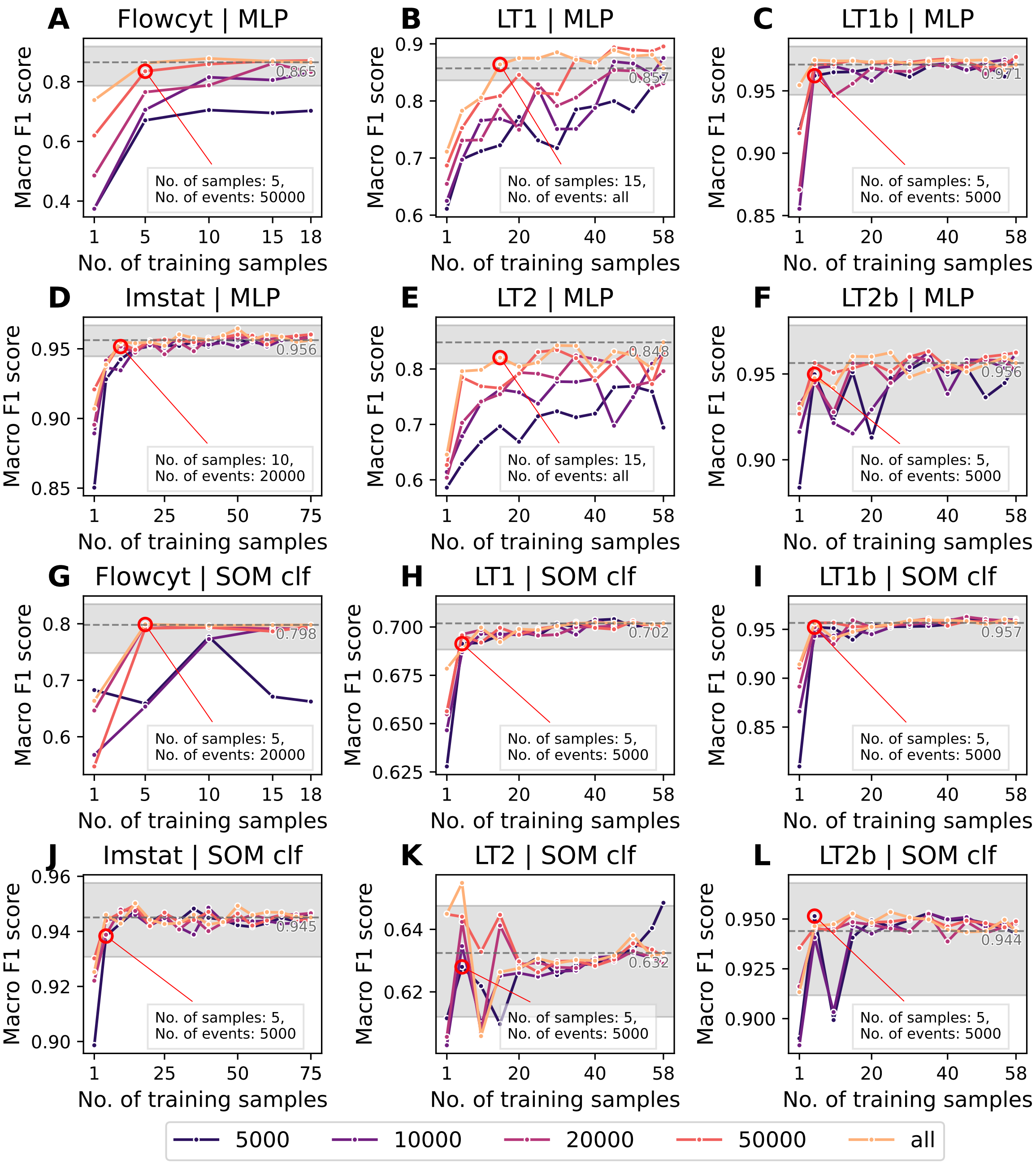


**Fig S7.** Average macro F1 scores across test samples of MLP and SOM classifier for varying numbers of training samples. Each curve corresponds to a different level of per-sample stratified downsampling, from 5000 events to all available events. Performance with the full training set is shown as a dashed line, shaded area around the line indicates the empirical 95 % confidence interval. Minimal number of samples ($> 1$) and minimal number of events for which performance lies within this confidence interval is highlighted with a red circle. These correspond to the minimal data requirements reported in Table 1 of the main manuscript. SOM classifier and MLP remain robust to reductions in sample count and events per sample, with performance improving quickly with few training samples and staying stable across downsampling levels. Even at extreme downsampling (5000 events $\lesssim$ 10% of events per sample), they maintain reasonable performance, indicating suitability for limited-data scenarios.

| Dataset | All Samples | All events |
|---|---|---|
| Flowcyt | 18 | 11671354 |
| Imstat | 75 | 2912192 |
| LT1 | 58 | 5413468 |
| LT2 | 58 | 5661119 |

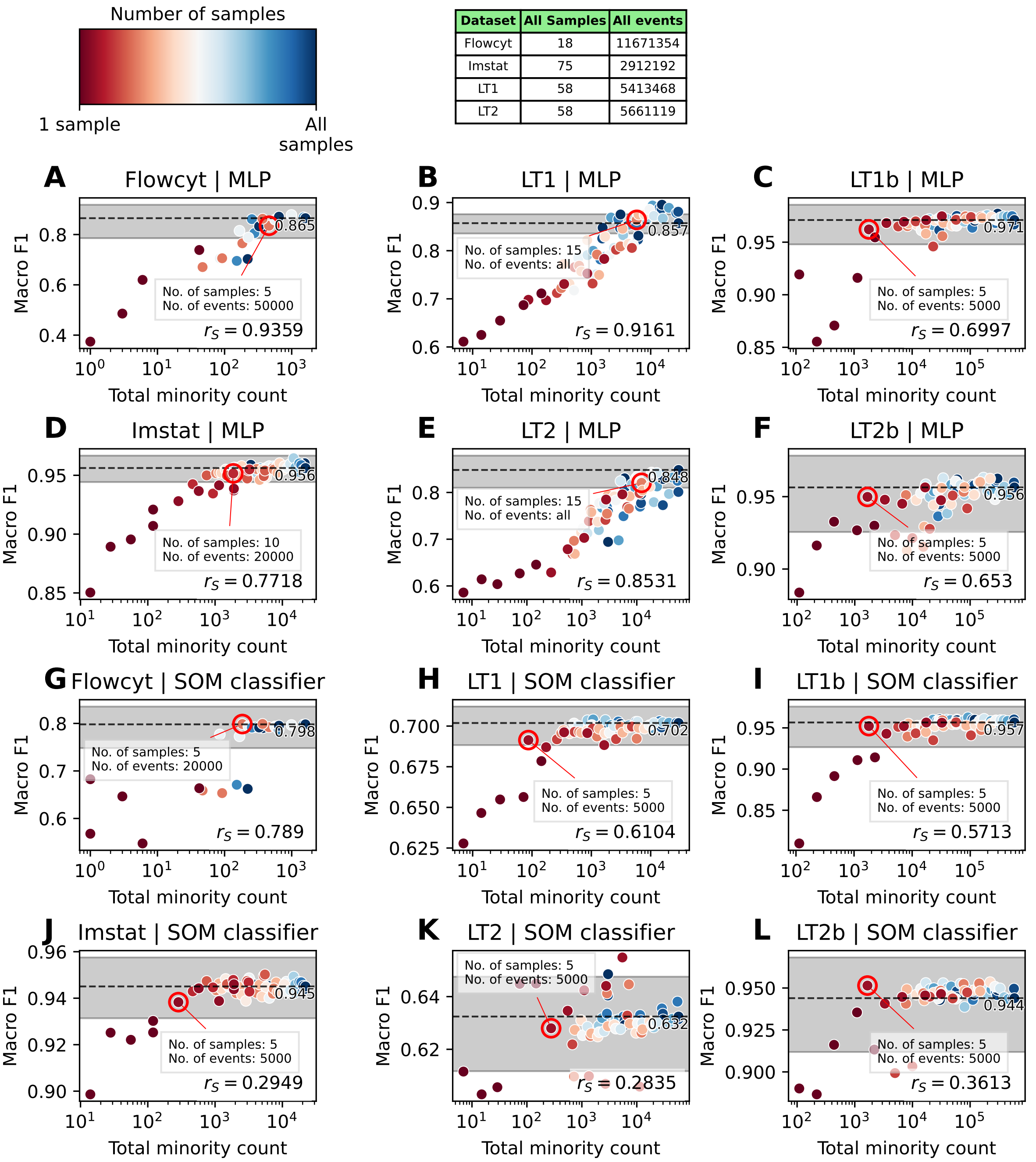


**Fig S8.** Average macro F1 scores across test samples against total number of events in the smallest class in the training samples (total minority count). Each point corresponds to a different level of per-sample stratified downsampling ($5 \times 10^3$, $10 \times 10^3$, $20 \times 10^3$, $50 \times 10^3$, all events) and a different number of samples included in the training set (1, 5, 10, …, all samples). Performance with the full training set is shown as a dashed line, shaded area around the line indicates the empirical 95 % confidence interval. Minimal number of samples ($> 1$) and minimal number of events for which performance lies within this confidence interval is highlighted with a red circle. These correspond to the minimal data requirements reported in Table 1 of the main manuscript. Spearman correlation is annotated in the lower right.

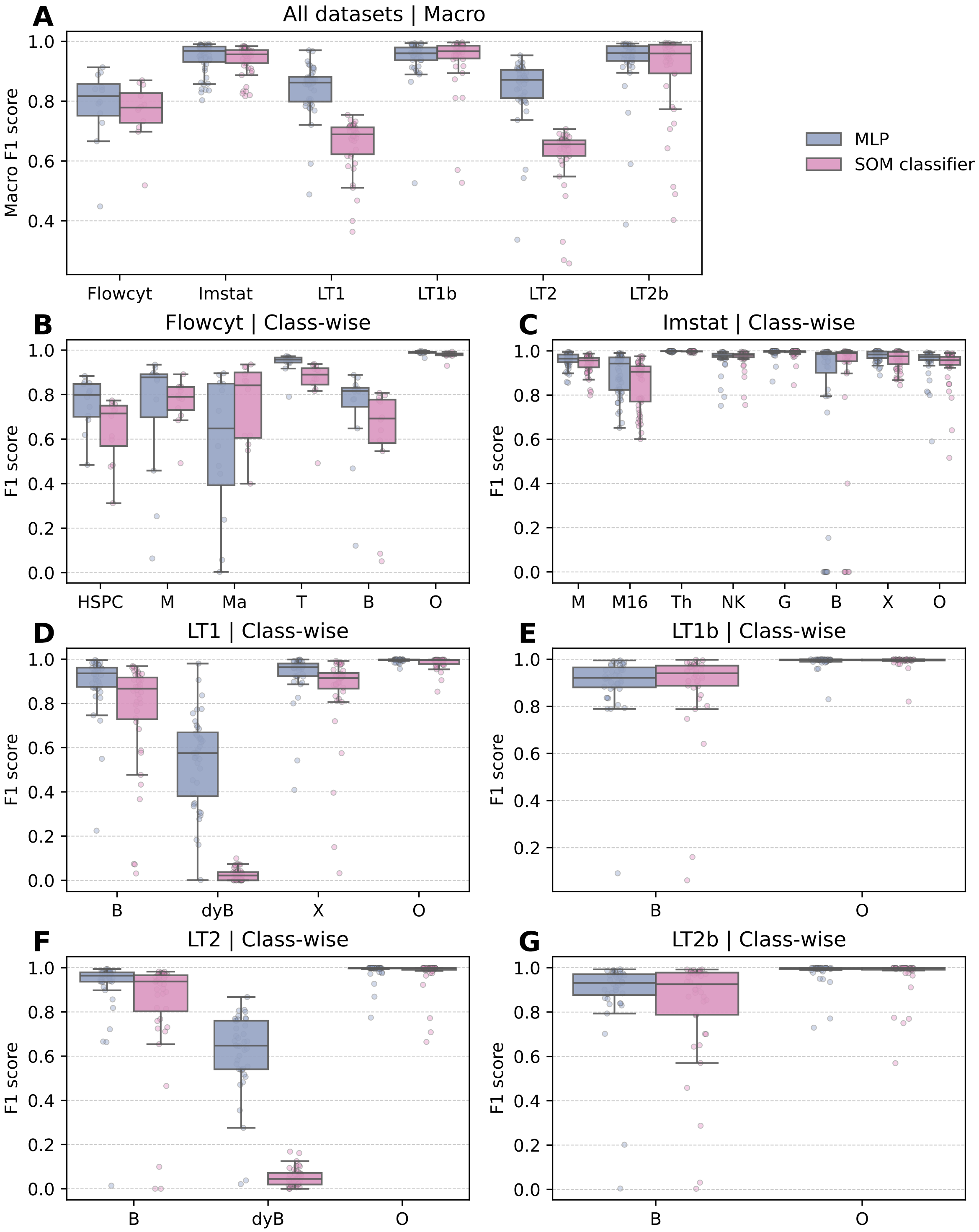


**Fig S9.** Macro and class-wise F1 scores across datasets for the locally trained MLP and SOM classier models. Training sets were randomly generated subsets of the respective complete training set with number of samples and number of events per sample set according to the minimal requirements reported in Table 2 of the main manuscript. Comparison to Fig S5 and Figure 1 of the main manuscript shows that models trained locally on reduced datasets achieve performance scores comparable to models trained on all samples.

### S4 Supplementary Tables

**Table S2.** LT1, LT1b. Letter to cell type label conversion and gating strategy.

| Integer Label | Full Name | Gating |
|---|---|---|
| B | B cells | CD19+/(+) and CD79b/CD4 +/(+) |
| dyB | (dying B cells) | B cells with low FS |
| X | debris, erythrocytes | CD45 negative |
| O | other cell types | all other cells |
| | **binary** | |
| B | B and dyB | |
| O | X and O | |

**Table S3.** LT2, LT2b. Letter to cell type label conversion and gating strategy.

| Letter Label | Full Name | Gating |
|---|---|---|
| B | B cells | SS low and (CD19+ or CD20+ or CD200+) |
| dyB | (dying B cells) | B cells and FS low |
| O | other cell types | all other cells |
| | **binary** | |
| B | B and dyB | |
| O | O | |

**Table S4.** Imstat. Letter to cell type label conversion. Gating was performed by one of the authors (MM) as described in [Plank et al., 2021]. M16 were gated as the CD14dim CD16+ subpopulation of monocytes.

| Letter Label | Full Name |
|---|---|
| B | B cells |
| Th | T helper cells |
| NK | Natural killer cells |
| M16 | Atypical, CD16 + monocytes |
| M | Monocytes, excluding M16 |
| G | Granulocytes |
| X | Outside the ususal gates for leukocytes, erythrocytes and debris, sorted out |
| O | Unclassified, other cells, i.e. the remainder (includes T killer cells, double negative T cells, basophils, precursor cells) |

**Table S5.** Flowcyt. Letter to cell type label conversion. Gating was performed by Bini et al. [2024].

| Letter Label | Full Name |
|---|---|
| T | T Lymphocytes |
| B | B Lymphocytes |
| M | Monocytes |
| Ma | Mast cells |
| HSPC | HSPCs |
| O | Others |

**Table S6.** Antibody marker channels used for gating and model training.

| Dataset | Antibody marker channels |
|---|---|
| Flowcyt | FS, SS, CD14, CD19, CD13, CD33, CD34, CD117, CD7, CD16, HLA, CD45 |
| Imstat | FS, SS, CD3, CD4, CD8, CD19, CD14, CD16, CD56, CD45 |
| LT1, LT1b | FS, SS, kappa+CD8, lambda+CD7, CD23, CD79b+CD4, CD5, CD38, CD19, CD20+CD3, FMC7+CD2, CD45 |
| LT2, LT2b | FS, SS, CD10, CD11c, CD19, CD20, CD25, CD43, CD52, CD103, CD200, IgM |

**Table S7.** Description of SOM classifier's hyperparameters and results of the first stage of the nested grid search.

| Parameter | Range | Description | Result |
|---|---|---|---|
| Grid topology | planar, toroid | topology of the SOM grid | planar is better |
| Grid type | rectangular, hexagonal | arrangement of SOM units in the 2D plane | rectangular is better |
| Grid dimensions | $(m,n) \in \mathbb{N} \times \mathbb{N}$ | dimensions of the SOM grid | moderate sizes work best ($m = n \in \{15, 20, 25\}$) |
| Neighborhood function | bubble, gaussian | function to compute neighborhood relations between SOM units | gaussian is better |
| $\sigma$ | $(0,\infty) \subset \mathbb{R}$ | parameter of the Gaussian neighborhood function | smaller is better ($\leq$ 1.0) |
| Initialization | random, PCA | initialization of weight vectors, at random or from the subspace spanned by the first two eigenvectors of the correlation matrix | PCA is better |
| Number of epochs | $T \in \mathbb{N}$ | number of passes of the training data | must be sufficiently large ($\geq 5000$) |
| Initial radius | $r_{\text{init}} \in (0,1] \subset \mathbb{R}$ | initial radius as a proportion of grid size $r(0) = r_{\text{init}} \cdot \min(m,n)$ | for $m = n$, values around $r_{\text{init}} = 0.5$ were best |
| Final radius | $r_{\text{final}} \in [0, \min(m,n)] \subset \mathbb{R}$ | - | smaller is better ($\leq$ 0.5) |
| Radius function | linear, exponential | parameterized such that such that $r(0) = r_{\text{init}} \cdot \min(m,n)$ and $r(T) = r_{\text{final}}$ | exponential is better |
| Initial learning rate | $\alpha_{\text{init}} \in (0,\infty) \subset \mathbb{R}$ | - | no clear pattern |
| Final learning rate | $\alpha_{\text{final}} \in [0,\infty) \subset \mathbb{R}$ | - | no clear pattern |
| Learning rate function | linear, exponential | parameterized such that such that $\alpha(0) = \alpha_{\text{init}}$ and $\alpha(T) = \alpha_{\text{final}}$ | exponential is better |

**Table S8.** Hyperparameter grid and best found value for SOM classifier.

| Parameter | Values in Grid Search | Optimal Value |
|---|---|---|
| Grid topology | planar | planar |
| Grid type | rectangular | rectangular |
| Grid dimensions | (15, 15), (20, 20), (25, 25) | (25, 25) |
| Neighborhood function | gaussian | gaussian |
| $\sigma$ | 0.1, 0.25, 1.0 | 0.1 |
| Initialization | PCA | PCA |
| Number of epochs | 1000 | 1000 |
| Initial radius | 0.25, 0.5, 0.75 | 0.25 |
| Final radius | 0.1 | 0.1 |
| Radius function | exponential | exponential |
| Initial learning rate | 0.1, 0.5, 1.0 | 0.1 |
| Final learning rate | 0.001, 0.05, 0.1 | 0.001 |
| Learning rate function | exponential | exponential |

**Table S9.** Computational environment used for the benchmark runs with all samples on NHR@FAU clusters. A single CPU was requested for GateMeClass, as its implementation is not parallelized.

| Job | Cluster | CPU Model | GPU | Allocated CPUs |
|---|---|---|---|---|
| GateMeClass | Woody (NHR@FAU) | 2 × Intel Xeon Gold 6326 (Ice Lake) | None | 1 |
| SOM classifier | Woody (NHR@FAU) | 2 × Intel Xeon Gold 6326 (Ice Lake) | None | 30 |
| MLP | TinyGPU (NHR@FAU) | 2 × Intel Xeon Gold 6134 (Skylake) | 1 × NVIDIA RTX 2080 Ti (11 GB) | 8 |
| DGCyTOF | TinyGPU (NHR@FAU) | 2 × Intel Xeon Gold 6134 (Skylake) | 1 × NVIDIA RTX 2080 Ti (11 GB) | 8 |

**Table S10.** Wall time and peak memory usage during model training on the full training set and inference on the test set. Durations longer than 10 seconds are rounded to the nearest second, while shorter durations are reported with two decimal places of precision. Memory usage values are reported as Gigabyte (GB) rounded to two decimals. for inference, values are reported as mean and standard deviation across individual test samples. Both DGCyTOF and MLP required 0.23 GB of peak GPU memory during training across all datasets. Note: Although DGCyTOF employs MLP as its base classifier, it trains faster because only 80 % of the data are used for training, with the remaining 20 % reserved as a validation set for confidence threshold calibration.

| **Dataset** | **Flowcyt** | **Imstat** | **LT1** | **LT2** | **LT1b** | **LT2b** |
|---|---|---|---|---|---|---|
| | | | **Training – Wall Time** | | | |
| GateMeClass | 1 h 13 m 35 s | 18 m 21 s | 28 m 31 s | 30 m 39 s | 25 m 37 s | 29 m 28 s |
| DGCyTOF | 1 h 22 m 10 s | 21 m 16 s | 39 m 40 s | 40 m 43 s | 38 m 48 s | 40 m 5 s |
| MLP | 1 h 44 m 48 s | 26 m 11 s | 49 m 3 s | 51 m 18 s | 48 m 53 s | 50 m 54 s |
| SOM-clf | 2 h 49 m 56 s | 1 h 9 m 21 s | 1 h 42 m 50 s | 1 h 46 m 30 s | 1 h 43 m 12 s | 1 h 46 m 2 s |
| | | | **Training – Peak Memory (CPU)** | | | |
| GateMeClass | 53.95 GB | 11.07 GB | 25.12 GB | 26.74 GB | 28.51 GB | 28.75 GB |
| DGCyTOF | 14.98 GB | 12.20 GB | 12.16 GB | 13.00 GB | 13.20 GB | 13.67 GB |
| MLP | 20.68 GB | 8.14 GB | 12.48 GB | 14.44 GB | 14.34 GB | 15.14 GB |
| SOM-clf | 61.52 GB | 15.73 GB | 28.87 GB | 30.71 GB | 29.28 GB | 30.59 GB |
| | | | **Inference – Wall Time**, (mean ± std) | | | |
| GateMeClass | 8 m 56 s ± 4 m 21 s | 4.33 ± 2.69 s | 11 ± 3.08 s | 13 ± 1.85 s | 11 ± 2.28 s | 11 ± 1.92 s |
| DGCyTOF | 4 m 24 s ± 1 m 6 s | 52 ± 12 s | 1 m 7 s ± 20 s | 52 ± 15 s | 58 ± 14 s | 55 ± 18 s |
| MLP | 0.58 ± 0.17 s | 0.03 ± 0.01 s | 0.07 ± 0.01 s | 0.05 ± 0.01 s | 0.05 ± 0.01 s | 0.06 ± 0.01 s |
| SOM-clf | 32 ± 9.84 s | 1.50 ± 0.65 s | 3.92 ± 0.49 s | 3.96 ± 0.57 s | 3.97 ± 0.57 s | 3.94 ± 0.52 s |
| | | | **Inference – Peak Memory (CPU)**, (mean ± std) | | | |
| GateMeClass | 45.71 ± 0.03 GB | 10.55 ± 0.00 GB | 23.74 ± 0.00 GB | 24.20 ± 0.55 GB | 27.85 GB ± 0.47 GB | 28.69 ± 0.00 GB |
| DGCyTOF | 11.39 ± 1.38 GB | 6.65 ± 1.05 GB | 6.96 ± 0.80 GB | 6.87 ± 0.62 GB | 6.91 ± 0.95 GB | 7.27 ± 1.45 GB |
| MLP | 4.41 ± 0.29 GB | 1.65 ± 0.03 GB | 2.32 ± 0.03 GB | 2.43 ± 0.02 GB | 2.40 ± 0.02 GB | 2.47 ± 0.02 GB |
| SOM-clf | 6.93 ± 1.22 GB | 1.34 ± 0.08 GB | 2.24 ± 0.06 GB | 2.45 ± 0.08 GB | 2.44 ± 0.06 GB | 2.49 ± 0.06 GB |
